# The origin and diversity of insulin sequences and structures in nature

**DOI:** 10.64898/2026.11.16.767061

**Authors:** Seyed Reza Moosavi, Zhenjian Lin, Eric W. Schmidt, Mark Yandell, Thomas Lund Koch, Helena Safavi-Hemami

**Author notes:** Corresponding author: Dr. Helena Safavi-Hemami.

## Abstract

The diversification of peptide signaling systems has been a major driver of metazoan evolution and biological complexity. The insulin superfamily, which includes insulins, insulin-like peptides, relaxins, and insulin-like growth factors, is an ancient group of peptide hormones that regulate development, growth, metabolism, and reproduction across Metazoa. Yet our understanding of the evolutionary origin of insulin signaling and the full extent of insulin superfamily diversity remain incomplete, particularly in invertebrates. Here, we performed a large-scale sequence and structural investigation of the insulin superfamily by mining public databases with sequence- and structure-based computational analyses to build a final dataset of 7,020 insulin-family sequences. Insulin-like peptides were present in all metazoan lineages except Porifera and Ctenophora. The absence of an identifiable canonical insulin ligand in Porifera is particularly intriguing because insulin receptors are present, raising questions about the receptor’s ancestral function and the evolutionary origin of its ligand. We further identified 12 distinct structural classes of insulin-superfamily peptides, with the greatest diversity in invertebrates, marking these lineages as a major source of insulin scaffold innovation. Together, these findings provide new insights into the origin and diversification of insulin signaling, establish a framework for understanding insulin superfamily evolution, and offer a resource for the discovery and engineering of therapeutic insulin analogs inspired by natural diversity.

## Introduction

Ancient signaling systems have persisted across animal evolution, while their components have been repeatedly modified, lost, and repurposed to support the remarkable diversity of animal forms, physiologies, and life histories. One such example is the insulin signaling system. Insulin is an ancient peptide hormone that plays key roles in metazoan biology, including development, growth, energy homeostasis, and metabolism (Le, et al. 2023). Insulin belongs to a broader superfamily of structurally related peptides, which includes insulin-like peptides (ILPs) in invertebrates and insulin, insulin-like growth factors (IGFs), and relaxins in vertebrates (Shabanpoor, et al. 2009).

Animals are broadly split into non-bilaterian lineages, including the early-diverging Porifera (sponges) and Cnidaria (jellyfish, corals, and sea anemones), and Bilateria, which includes the vast majority of animal species and is further divided into protostomes (e.g., arthropods and mollusks) and deuterostomes (e.g., echinoderms and vertebrates). The earliest animal lineage known to possess an insulin-like peptide ligand and insulin receptor is cnidarians (Steele, et al. 1996; Anctil 2009), a lineage that diverged from bilaterians more than 600 million years ago (Kraus, et al. 2016). In Porifera, an earlier-diverging phylum, putative components of the insulin signaling system have been suggested, however, available evidence remains limited and inconclusive (Robitzki, et al. 1989; Skorokhod, et al. 1999). Thus, the evolutionary origin of the insulin signaling system remains unresolved.

The origin of a distinct relaxin signaling system is likewise incompletely understood, although relaxin-like peptides have been described across multiple protostome phyla suggesting an early bilaterian origin prior to the protostome-deuterostome split (Feng, et al. 2025). For example, in *Drosophila melanogaster*, the insulin-like peptide Dilp8 has relaxin-like activity and acts as a ligand for a relaxin-like receptor (Garelli, et al. 2015; Vallejo, et al. 2015), suggesting the presence of a relaxin-like system in protostomes. In contrast, the emergence of a distinct IGF signaling system has been regarded as a chordate innovation. This hypothesis is supported by the discovery of a single insulin-like peptide in the cephalochordate amphioxus (*Branchiostoma* spp.) that possesses structural features of both insulin and IGF, suggesting that it resembles the ancestral precursor from which both the insulin and IGF systems subsequently diverged (Chan, et al. 1990). The earliest evidence for three distinct insulin superfamily genes with similarity to insulin, IGF, and relaxin comes from the vase tunicate *Ciona intestinalis* (Olinski, et al. 2006), an ascidian species belonging to the subphylum Urochordata, the closest living relatives of vertebrates (Delsuc, et al. 2006). Together, these findings support a model in which an ancestral insulin signaling system emerged early in non-bilaterians and diversified progressively into a multi-member peptide superfamily during metazoan evolution.

All insulin superfamily members are synthesized as prepropeptides that are processed into mature peptides adopting a conserved “insulin fold” that is critical for biological activity (Adams, et al. 1969; Rao, et al. 2024). In vertebrates, mature insulin consists of two peptide chains (A and B chain) connected by two interchain disulfide bonds, with an additional intra-chain disulfide bond in the A chain. Insulin is initially translated with a connecting C-peptide, which is proteolytically removed after folding and disulfide bond formation to yield the mature two-chain hormone (Weiss, et al. 2000). Relaxins share the chain architecture and disulfide bonding pattern. In contrast, IGFs are single-chain peptides that retain the C-peptide region and include an additional D domain at the C-terminal (Weiss, et al. 2000).

Insulins, IGFs, and likely most ILPs signal through related receptor tyrosine kinases, in humans most prominently through the insulin receptor (IR) and IGF-1 receptor (IGF1R), which activate downstream signaling pathways such as PI3K–Akt and MAPK (Avruch 1998; Taniguchi, et al. 2006). Functionally, insulin signaling predominantly regulates metabolic homeostasis, including glucose uptake, whereas IGF signaling is more closely associated with cell proliferation and developmental programs (De Meyts 1994; Jensen, et al. 2007). In contrast, relaxins signal through G protein-coupled receptors (GPCRs) of the relaxin family and regulate reproduction (Feng, et al. 2025) and metabolism (Bathgate, et al. 2013).

Although less studied than their vertebrate homologs, invertebrate ILPs exhibit more structural variation, including differences in chain lengths, disulfide connectivity, and posttranslational modifications. For example, the genome of the nematode *Caenorhabditis elegans* encodes 40 ILPs that can be classified into multiple structural classes (Pierce, et al. 2001), while the genome of the Pacific oyster, *Crassostrea gigas*, encodes six ILPs with distinct disulfide scaffolds (Cherif-Feildel, et al. 2019). A mature ILP from the sea hare, *Aplysia californica*, was shown to contain multiple posttranslational modifications (Floyd, et al. 1999). Insects also display diverse sequences and structures with 8 ILP genes in the fruit fly *D. melanogaster* (Wu and Brown 2006) and more than 30 genes in the domestic silk moth, *Bombyx mori* (Aslam, et al. 2011). Despite this diversity, structural characterizations remain limited, with only a small number of ILPs from *C. elegans* and *D. melanogaster* assessed to date (Mizoguchi and Okamoto 2013).

In vertebrates, disruption of insulin superfamily signaling is associated with various diseases, most prominently, diabetes mellitus (DM) (Bathgate, et al. 2013; White and Kahn 2021; Van de Weyer and Tahas 2024; Divers 2026). Insulin is essential for regulating glucose homeostasis and remains a life-saving therapy for millions of people with DM. However, the native human insulin molecule has physicochemical properties that limit its therapeutic utility, which continues to spark the development of novel insulin analogs with improved properties (Perkins, et al. 2021). Notably, several analogs have drawn inspiration from naturally occurring insulins. For example, guinea pig insulin, which is monomeric, has been explored as a fast-acting insulin (Zimmerman and Yip 1974; Engholm, et al. 2015), while avian variants, such as chicken and turkey (Simon, et al. 1974; Muggeo, et al. 1979) exhibit increased potency at the human IR. Beyond vertebrates, monomeric ILPs from cone snail venom have been investigated as templates for ultra-fast-acting insulin (Safavi-Hemami, et al. 2015; Xiong, et al. 2020; Xiong, et al. 2022), and viral insulins from the Iridoviridae family have provided tools to study insulin receptor agonism and antagonism (Altindis, et al. 2018; Zhang, et al. 2021). Beyond this primary structure, divergent insulin scaffolds that contain additional disulfide bonds to enhance insulin’s limited stability have also been explored (Wu, et al. 2017). Together, these findings illustrate how the structural and pharmacological diversification of the insulin superfamily throughout metazoan evolution has generated a rich repertoire of naturally occurring molecules that can be leveraged as templates for the design of next-generation insulin analogs with improved properties. However, the full extent of this diversity remains largely unexplored, particularly outside vertebrates, highlighting the need for systematic efforts to uncover insulin-family peptides across diverse metazoan phyla.

Several recent studies have begun to explore insulin peptide diversity in animals, including large-scale surveys spanning major lineages (Chowański, et al. 2021; Irwin 2021; Rao, et al. 2024; Chrudinová, et al. 2026). While these studies have contributed greatly to our understanding of insulin family evolution, these efforts did not systematically address the evolutionary origin of the insulin signaling system or fully explored the extent of sequence and structural diversity within the superfamily, particularly in protostomes. Here, by mining publicly available protein sequence databases and using cysteine scaffold arrangements to identify sequence outliers with potentially unusual folds, we present the most comprehensive sequence and structural survey of the insulin superfamily to date. We identify insulin-like sequences across nearly all animal phyla, except Ctenophora and Porifera, provide new insights into the origin and distribution of the insulin signaling system, and reveal a diverse array of previously unrecognized structural scaffolds. Coevolutionary analyses further reveal residue-level constraints governing insulin folding, offering a framework for the rational design of next-generation therapeutic insulins. This dataset, comprising 7,020 sequences, represents a valuable resource for studying lineage-specific insulin evolution and for leveraging natural diversity in future analog design.

## Results

### Iterative sequence mining expands the known repertoire of the insulin superfamily

To systematically map the sequence space of the insulin superfamily, we began with 243 validated seed sequences spanning the major insulin-family groups from Feng et al. (Feng, et al. 2025), and employed iterative PSI-BLAST searches against the NCBI non-redundant (NR) database (Supporting Figure 1A). After removing duplicates, this procedure yielded ∼18,000 sequences. We removed duplicate, chimeric, non-target, and sequences shorter than 50 or longer than 500 amino acids, then clustered the remaining filtered sequences at 95% identity, retaining 7,200 sequences. These sequences were folded *in silico* using ESMFold (Lin, et al. 2023), and we manually inspected the longest and shortest examples to confirm the presence of the insulin fold. Following these filtering steps, the dataset contained 6,969 high-confidence insulin-family sequences.

**Figure 1.**
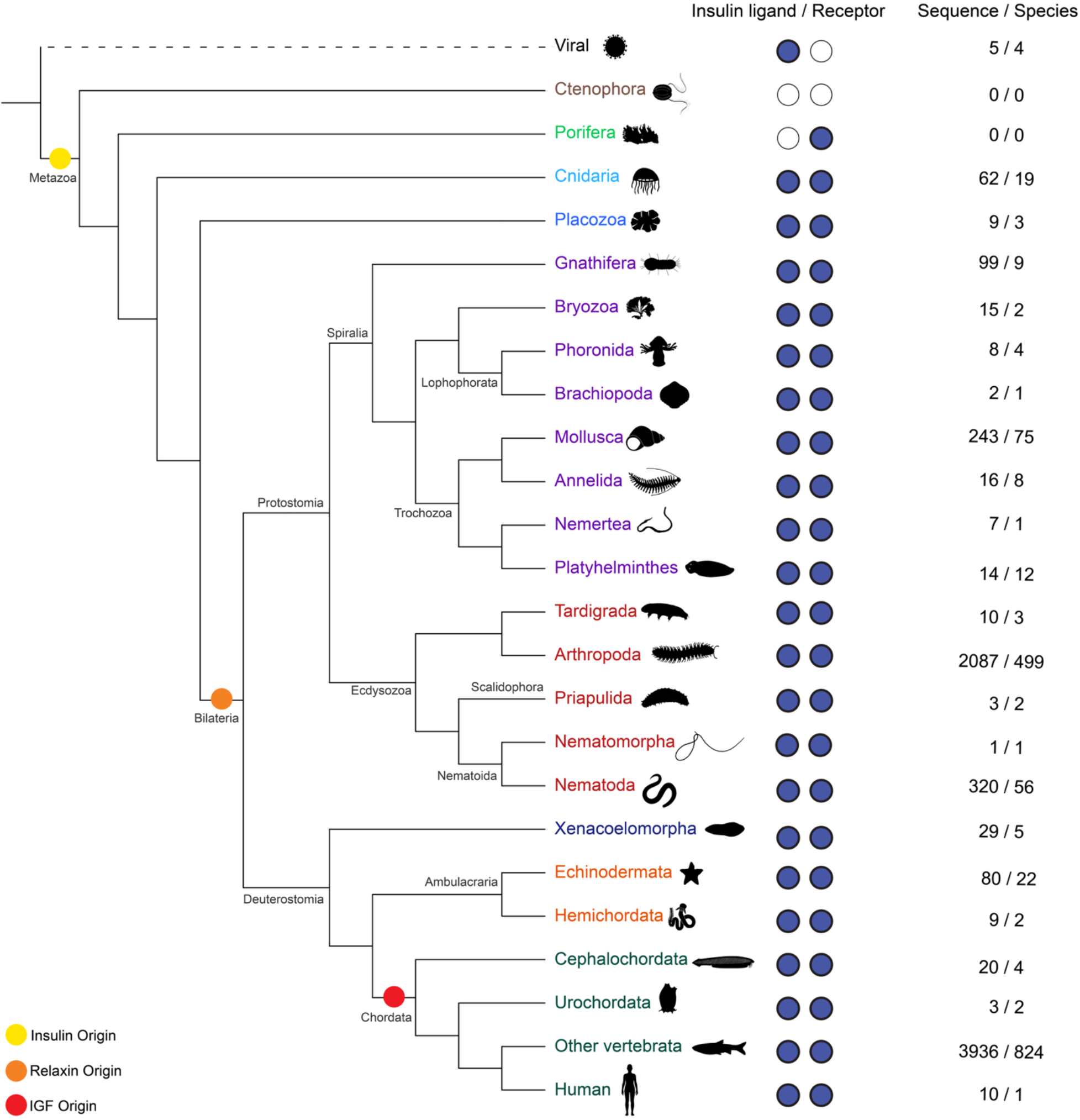
Cladogram showing the phylogenetic distribution of insulin superfamily sequences across Metazoa, highlighting near-ubiquitous presence and broad taxonomic coverage of the dataset (7,020 sequences). Each phylum or major lineage is annotated with the total number of sequences and represented species, and the presence or absence of insulin ligands and insulin receptors is indicated by filled or empty circles, respectively. Bilaterian lineages dominate the dataset, with particularly large representation in Chordata, Arthropoda, and Nematoda, whereas early-branching lineages such as Cnidaria, Placozoa, and Xenacoelomorpha are sparsely represented. No insulin-superfamily sequences were detected in Porifera or Ctenophora. Viral insulin-like peptides are displayed as a separate outgroup due to their distinct evolutionary origins. Colors denote major metazoan clades. Putative origins of insulin, relaxin, and IGF are indicated by circles. Organism silhouettes in Figure 1 were obtained from PhyloPic (https://www.phylopic.org) and are used under their respective licenses.

Sequences from some metazoan phyla were absent from our dataset, likely because of greater sequence divergence or limited representation in the NR database. For these phyla (Placozoa, Phoronida, Xenacoelomorpha, Platyhelminthes, and Nemertea), we added sequences reported in the literature (Nikitin 2015; Thiel, et al. 2021) to our dataset and conducted targeted homology searches of transcriptome shotgun assemblies (TSAs) from representative species. This identified additional insulin sequences from these phyla, yielding a final dataset of 7,020 sequences used for downstream analyses (Supporting File 1). This dataset represents the most comprehensive collection of insulin superfamily sequences compiled to date.

### Insulins are present across metazoan phyla except Porifera and Ctenophora

We used this curated database to characterize the distribution of insulin-family sequences across metazoan phyla, which revealed a broad and nearly ubiquitous presence (**Figure 1**). Given species biodiversity and species bias in the NR-database, sequences derived from bilaterians dominate the dataset, with a particularly large number of sequences recovered from chordates (3,969 sequences from 831 species) and major protostome groups such as Arthropoda (2,087 sequences from 499 species) and Nematoda (320 sequences from 56 species) (**Figure 1**). In contrast, early-branching metazoan phyla were sparsely represented. Cnidaria and Placozoa contained only 62 and 9 sequences, respectively, while no sequences were detected in Porifera or Ctenophora. Insulins reported from viruses of the Iridoviridae family were also retrieved, although their evolutionary origin remains unresolved (Supporting File 1).

The apparent absence of insulin-like sequences in Ctenophora has previously been reported (Sachkova, et al. 2021; Hayakawa, et al. 2022). Consistent with this, we failed to identify any insulin-like peptides in ctenophores, either in our initial searches or in additional analyses incorporating both sequence-and structure-based interrogation of published ctenophore sequencing datasets. These findings support the hypothesis that the insulin signaling system emerged after the divergence of Ctenophora.

Our initial searches also revealed no hits in Porifera. However, given reports of an insulin-like peptide in the sponge *Geodia cydonium* (Robitzki, et al. 1989) and insulin-like receptors in several sponge species (Suzuki, et al. 2000), we assessed transcriptomic data from 220 sponge species for the presence of putative insulin-like peptides. Using BLASTp searches with diverse insulin-like sequences as queries, hidden Markov models (HMMs) (Johnson, et al. 2010), and structure-based searches (van Kempen, et al. 2022), insulin-like sequences were only detected in 64 of 1,945 transcriptome samples (corresponding to 37 of 220 species). A species-by-sample heatmap revealed an irregular within-species distribution, with hit-positive libraries scattered among predominantly negative libraries rather than being consistently recovered across conspecific transcriptomes (Supporting Figure 2). Independent screening against a panel of sponge genomes and genome-derived proteomes showed that none of the insulin hits mapped to the genome assemblies. Taken together, this pattern is difficult to reconcile with the presence of a conserved bona fide sponge insulin gene and instead strongly suggests sample contamination in cases where insulin-like sequences were recovered. The previously reported cDNA sequence from *Geodia cydonium* (Robitzki, et al. 1989) was also determined to be a contaminant, as it shares > 80% identity with insulin sequences from multiple rodent species and has since been re-annotated as deriving from *Rodentia sp.* (taxon ID: 69158, NCBI Sequence ID: P21563.1, Supporting File 1).

**Figure 2.**
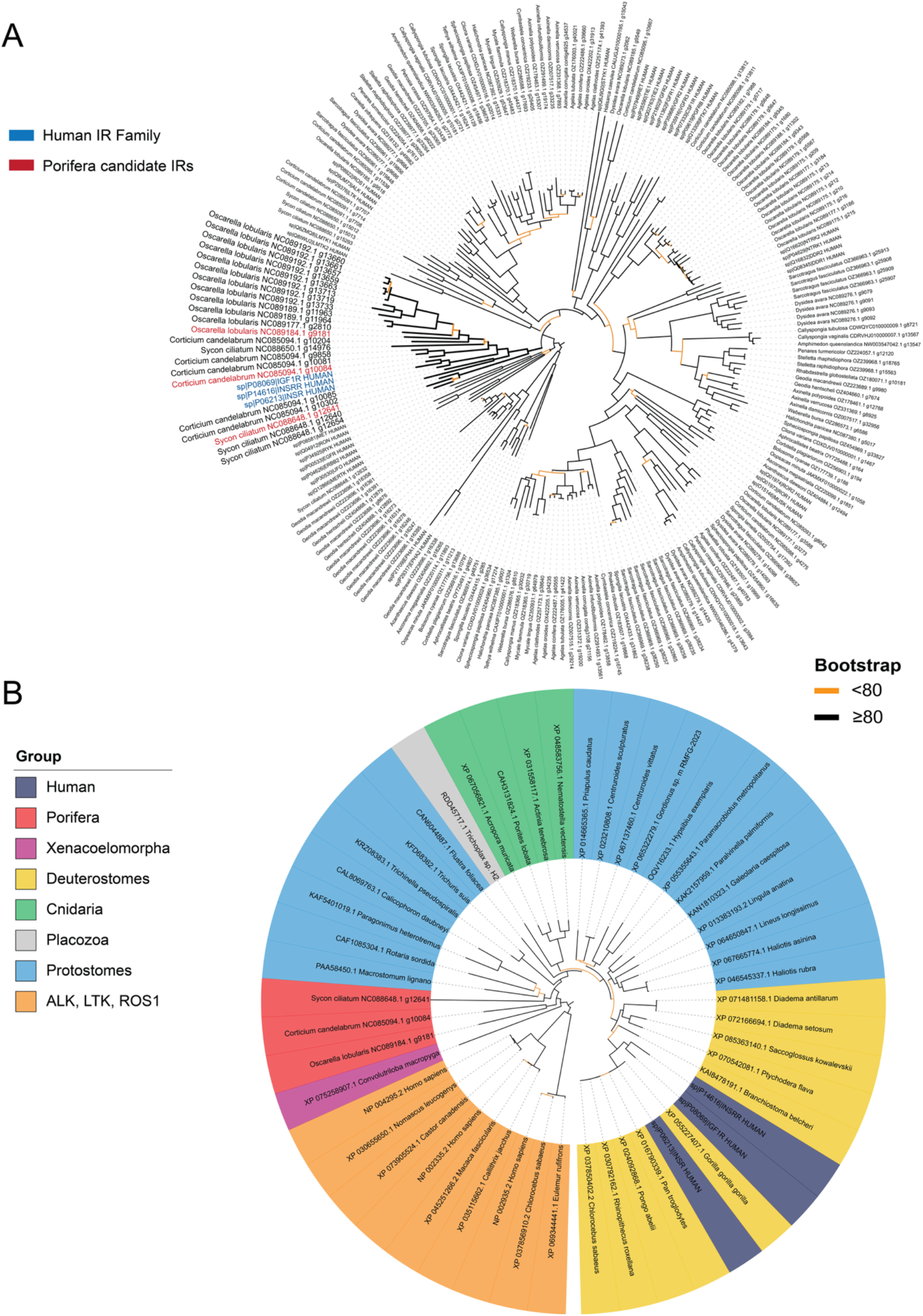
Poriferan insulin receptors form an early-diverging lineage within the animal insulin receptor family. A) Maximum-likelihood phylogenetic analysis of 187 poriferan candidate sequences with confirmed genomic locations, together with the complete set of human receptor tyrosine kinases (RTKs), identified sequences from *Oscarella lobularis*, *Corticium candelabrum*, and *Sycon ciliatum* that group with the human insulin receptor family rather than with other RTK families. The human insulin receptor family is indicated in blue, and the poriferan sequences selected for further analysis are highlighted in red. B) Phylogenetic analysis of representative insulin receptors from major animal lineages places the poriferan receptors in a distinct lineage sister to the insulin receptors of all other sampled animals. This position is consistent with their origin in Porifera and supports their identification as genuine insulin receptors. ALK, LTK, and ROS1 sequences form the outgroup to the insulin receptor clade. Taxonomic groups are indicated by the colored sectors. Branches with bootstrap support ≥80 are shown in black, whereas branches with support <80 are shown in orange.

### Insulin receptors are not detected in Ctenophora but present in Porifera

The apparent absence of insulin-like sequences in Ctenophora and Porifera prompted us to search for homologs of the insulin receptor (IR) in these lineages to determine if the signaling system is lacking or if components of the system are already found in these early divergent lineages. Consistent with the apparent absence of an insulin-like peptide in Ctenophora, BLASTp searches against the NCBI NR database using the human IR as query failed to identify any similar sequences in this phylum. In contrast, several sponge species have previously been reported to express putative IR-like proteins (Skorokhod, et al. 1999), and BLASTp searches against the NCBI NR and sponge transcriptome datasets readily identified multiple IR-like sequences (Supporting File 1). To determine whether these sequences belong to the IR family rather than with other receptor tyrosine kinases (RTKs), we constructed a maximum-likelihood phylogeny combining 187 Porifera candidate IR-like sequences from our in-house database, each with a confirmed genomic location, with the complete set of human RTKs categorized by Filis et al. (Filis, et al. 2023). Of the poriferan species analyzed, only sequences from *Oscarella lobularis* and *Corticium candelabrum*, both members of Homoscleromorpha, and *Sycon ciliatum*, a member of Calcarea, clustered with the human IR family; the remaining candidates grouped with other RTK families (**Figure 2A**; Supporting File 2). Thus, IR homologs were identified in two of the four major classes of Porifera.

To further evaluate the identity and evolutionary placement of the poriferan IR sequences, we constructed a second maximum-likelihood phylogeny containing representative IRs from the major animal phyla shown in Figure 1. Three representative sequences from each of the closely related RTK families ALK (anaplastic lymphoma kinase), LTK (leukocyte tyrosine kinase), and ROS1 (ROS proto-oncogene 1) were included as outgroups. In this broader analysis, the poriferan sequences formed a distinct lineage associated with the IR clade and were positioned outside the protostome and deuterostome IR groups (**Figure 2B**; Supporting File 2). This placement is consistent with the early divergence of Porifera and argues against contamination from protostome or deuterostome species.

To further assess their origin, we examined the genomic context of the candidate IR-like genes in *C. candelabrum* and *O. lobularis*. In both species, the IR-like genes localize to Chromosome 10, providing additional evidence that they are genuine poriferan genes rather than contaminants (Supporting File 2). Together, these findings support a model in which the insulin signaling system emerged after the divergence of Ctenophora, with the receptor component already present in Porifera and the canonical insulin–insulin receptor signaling pair established in Cnidaria. Whether an insulin-like peptide was present in Porifera and subsequently lost, or instead arose later in animal evolution, remains unresolved.

### Insulin superfamily phylogeny highlights canonical protostome–deuterostome divergence with some exceptions

To resolve evolutionary relationships across the insulin superfamily, we constructed a maximum-likelihood phylogenetic tree of the 7,020 sequences in our database (**Figure 3**, Supporting File 3). Sequences were labeled according to one of five major lineages (Cnidaria, Placozoa, Protostomia, Xenacoelomorpha, and Deuterostomia) based on taxonomic information extracted from sequence headers using CLANS, with sequences retrieved from the literature labeled manually. Phylogenetic analyses showed that sequences broadly resolved based on their taxonomic lineage, with protostome insulin-like peptides (ILPs) forming two major groups and deuterostome sequences resolving into three groups corresponding to insulins, IGFs, and relaxins based on annotated reference sequences. Cnidarian, placozoan, and Xenacoelomorpha sequences did not form independent clades but instead nested within the protostome ILP group. Furthermore, consistent with their reported sequence similarity to IGFs, viral insulin-like peptides group within IGFs.

**Figure 3.**
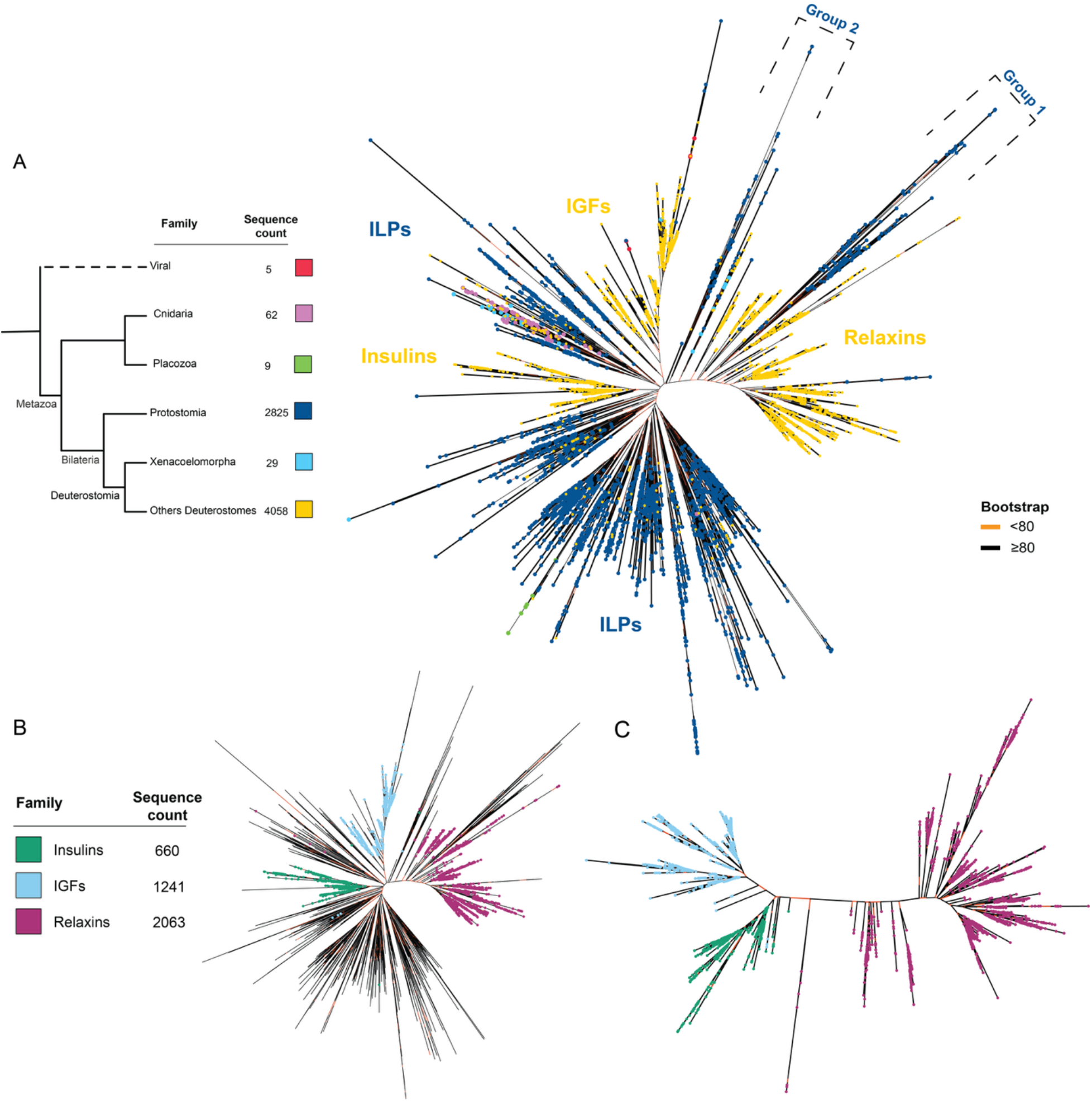
Maximum-likelihood (ML) phylogenetic analyses reveal evolutionary structure within the insulin superfamily. A) Full-dataset phylogeny of 7,020 insulin-family sequences, colored by phylum-level classification: Deuterostomia (yellow), Protostomia (dark blue), Placozoa (green), Xenacoelomorpha (blue), Cnidaria (purple), and viral sequences (red). The tree was inferred with IQ-TREE2 under the best-fit substitution model selected by ModelFinder Plus. Bootstrap support values are displayed on the trees, with orange indicating support <80 and black indicating support ≥80. Locations of minor phylum-level groups are annotated, and two divergent protostome clades, Group 1 and Group 2, are highlighted. B) The same full tree recolored by insulin-superfamily subgroup (insulin, IGF, relaxin) based on CLANS cluster-based sequence annotations as shown in Figure 4A. C) A focused ML phylogeny containing only deuterostome insulin, IGF, and relaxin sequences. Bootstrap support values are shown using the same color scheme as in panel A (orange <80; black ≥80).

Protostome and deuterostome sequences largely segregated into different groups, with two notable exceptions. First, the deuterostome insulin clade fell between the two protostome ILP groups rather than clustering with deuterostome IGFs and relaxins. Second, two small clades of protostome ILPs (Group 1 and Group 2) nested within the deuterostome relaxin clade, raising the possibility that these represent protostome members of the relaxin family (**Figure 3**, Supporting Figure 3, Supporting File 3). Since Drosophila Dilp8 has previously been shown to be relaxin-like (Garelli, et al. 2015; Vallejo, et al. 2015), we included Dilp8-like sequences as marker of these two divergent protostome clades, alongside canonical deuterostome relaxins, insulins, and IGFs, in a phylogenetic analysis of this subset (Supporting Figure 3). Most Group 2 sequences, along with a subset of Group 1, clustered with Dilp8-like peptides, while the remaining Group 1 sequences clustered with canonical relaxins rather than with insulin-or IGF-like sequences. Together, these results support classifying both Group 1 and Group 2 as relaxin-like peptides, with Group 2 showing closer affinity to the protostome Dilp8-like lineage and Group 1 more closely resembling canonical relaxins. This is consistent with the broader topology observed in Figure 3A-B.

Highlighting of annotated insulins, IGFs, and relaxins (**Figure 3B**) confirmed that the three families largely occupy distinct regions of the tree, consistent with their classification as separate deuterostome lineages within the insulin superfamily. An independent phylogenetic tree using only these annotated deuterostome insulin, IGF, and relaxin sequences (660, 1,241, and 2,063 sequences, respectively) further resolved the three families into distinct clades (**Figure 3C**).

### CLANS and protein language model analyses independently confirm insulin superfamily family- and lineage-level structure

As phylogenetic analyses can provide limited resolution, particularly for short and divergent sequences, we complemented our tree-based approach with CLANS cluster analysis (Frickey and Lupas 2004), which uses all-versus-all BLASTp similarity scores to organize sequences into similarity-based clusters independent of tree topology (Supporting File 4). Deuterostome sequences formed three distinct clusters, which could be classified as insulins, IGFs, and relaxins based on the presence of well-annotated reference sequences within each cluster (**Figure 4A**, dashed circles). In contrast, protostome sequences resolved into numerous smaller, more fragmented clusters. Sequences from Cnidaria and Placozoa each formed distinct clusters, whereas Xenacoelomorpha sequences were more broadly dispersed across the map. A more detailed breakdown of these deeper taxonomic groupings is provided in Supporting File 4. This clustering pattern closely mirrored the topology of the phylogenetic tree, with deuterostome sequences again resolving cleanly into insulin, IGF, and relaxin subfamilies while protostome sequences formed a more fragmented, less clearly resolved structure. To provide another independent line of support for the separation of protostome and deuterostome sequences, as well as the distinct clustering of insulin, IGF, and relaxin families, we generated sequence embeddings for all insulin superfamily members using the protein language model ESM-2 (Lin, et al. 2023), followed by dimensionality reduction via UMAP (Supporting File 4). This alignment-free approach captures sequence similarity based on learned biochemical and structural features rather than explicit sequence alignment, providing a means of validation independent of both phylogenetic and CLANS-based analyses. The resulting embeddings largely recapitulated the clustering patterns observed by CLANS (**Figure 4B**): sequences segregated into clearly defined groups corresponding to protostome and deuterostome lineages, with non-bilaterian ILPs again grouping among protostome sequences. To assess whether subfamily-level structure was similarly recoverable, we next colored the UMAP embedding according to the insulin, IGF, and relaxin groupings identified in Figure 4A (Supporting Figure 4). The three deuterostome subfamilies formed distinct, well-separated clusters in embedding space, mirroring their separation in the phylogenetic tree and confirming that subfamily-level divergence within the insulin superfamily is recoverable from sequence information alone, independent of alignment or phylogenetic inference.

**Figure 4.**
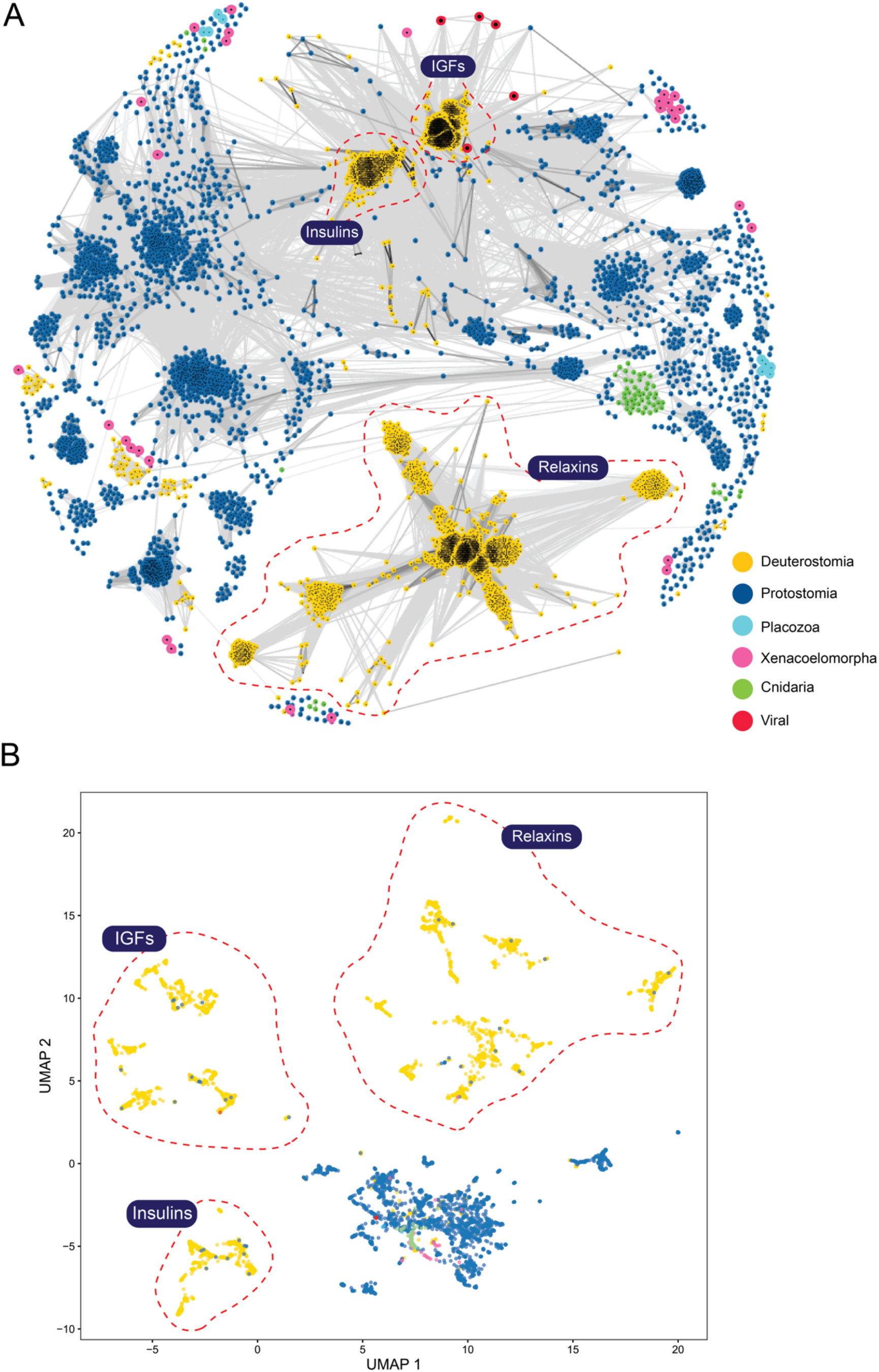
CLANS clustering of insulin superfamily sequences reveals distinct deuterostome clusters corresponding to the major insulin families, contrasted with more diverse and diffuse protostome groupings. A) Two-dimensional CLANS map generated from all-against-all BLASTp comparisons of the curated dataset of 7,020 insulin superfamily sequences. Each point represents an individual sequence, colored by phylum-level taxonomy (Deuterostomia, Protostomia, Placozoa, Xenacoelomorpha, Cnidaria, and viral sequences). Attraction–repulsion forces position sequences according to pairwise similarity. Three densely populated regions enriched for well-annotated representatives of insulin, IGF, and relaxin-family peptides are marked by dashed circles. Points representing sequences from Placozoa, Xenacoelomorpha, and viral sequences were enlarged using Adobe Illustrator for improved visibility. Gray connecting lines (p-value threshold ≤1×10⁻⁷) indicate high similarities between sequences. B) Clustering of sequences using embedding vectors from the ESM-2 protein language model, using the same color scheme as in panel A. ESM-2-based clustering largely recapitulates the CLANS results, recovering the deuterostome-protostome/non-bilateria separation as well as the grouping of insulin-family members (insulin, relaxin, and IGF).

### The insulin superfamily consists of multiple structural classes

Our phylogenetic analyses support an ancient origin of the insulin signaling system followed by extensive diversification of insulin-family peptides throughout metazoan evolution. In addition to supporting this evolutionary framework, our systematic mining of publicly available sequence databases generated a comprehensive resource of ILP, insulin, IGF, and relaxin homologs spanning nearly every major animal phylum. We next leveraged this resource to investigate whether this diversification has generated previously unrecognized cysteine architectures and, potentially, novel structural scaffolds, diversity that could both expand our understanding of insulin-family biology and provide new templates for next-generation insulin analog design.

We first attempted to cluster predicted three-dimensional structures generated by ESMFold (Lin, et al. 2023) (Supporting File 5) using Foldseek (van Kempen, et al. 2022). This approach proved ineffective, as clustering was dominated by variation in flexible, disordered regions such as the C-peptide rather than differences within the conserved insulin core, and structure predictions themselves represent computational hypotheses of uncertain accuracy. We therefore developed a classification strategy based on cysteine patterns, which are determined directly from primary sequence rather than inferred structure. Because cysteine number and position encode the potential disulfide-bonding architecture of these peptides, substantial changes in either provide direct, sequence-level evidence of departure from the canonical insulin scaffold, independent of whether the precise three-dimensional consequences can be confidently predicted. Briefly, cysteine positions were extracted from signal-peptide-trimmed sequences, mapped to their corresponding multiple sequence alignment (MSA) column indices, and used to group sequences by shared connectivity pattern (Supporting Figure 5). One representative sequence per group was retained for manual inspection of sequence and predicted structure.

**Figure 5.**
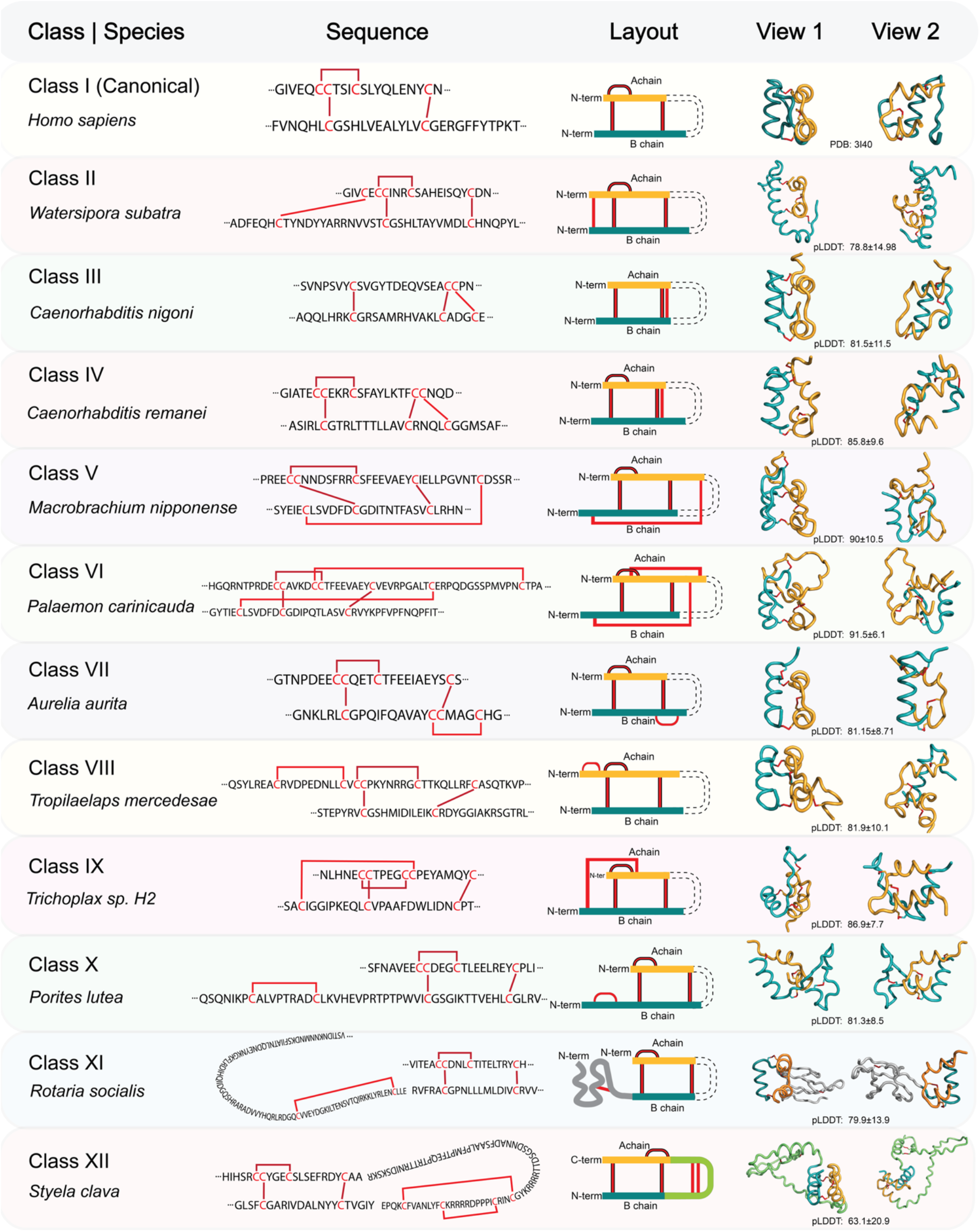
Representative insulin-family peptides from twelve cysteine-pattern classes (Class I–XII) identified across metazoan lineages, highlighting the diversity of sequence-defined cysteine frameworks within the insulin superfamily. For each class, a representative species, sequence, cysteine-layout schematic, and AlphaFold3-predicted structure (shown in two views) are provided. A and B chains are shown in orange and teal, respectively. The twelve classes are defined by the number and relative positions of cysteine residues in the sequence; thus, class assignment is based on sequence features rather than on experimentally determined disulfide connectivity or three-dimensional structure. Predicted structures, together with AlphaFold confidence scores, are provided as Supporting File 5. Disulfide bonds shown in red in the structural models represent predicted cysteine connectivity and should not be interpreted as experimentally validated. Canonical Class I cysteine positions are outlined with black borders to distinguish them from additional or alternative cysteine patterns. Boundaries between A- and B-chain regions are approximate and are not based on experimentally validated or computationally predicted cleavage sites. This uncertainty in chain boundaries and processing is indicated by three-dot markers at the chain termini. All sequences depicted in the figure, together with additional sequences representing each class, are provided in Supporting File 5.

This analysis identified 12 distinct cysteine-pattern classes, each defined by a characteristic disulfide-bonding architecture (Supporting File 5: PSE Files; **Figure 5**). We used AlphaFold3 to predict the structure of a representative sequence from each class (Supporting File 5), calculating mean per-residue confidence (pLDDT) across the core fold containing the cysteine residues and their disulfide connectivity, while excluding signal peptides, C-peptides, and extended terminal regions (script provided in Supporting File 7). These structures should be interpreted as predictions rather than experimentally validated models; critically, however, the classification itself is independent of their accuracy, as it rests on sequence-encoded cysteine patterns rather than predicted folds. While the precise three-dimensional consequences of individual classes will require experimental validation, their distinct cysteine architectures provide strong sequence-level evidence for substantial structural diversification within the insulin superfamily.

Class I, exemplified by human insulin, corresponds to the canonical insulin cysteine architecture that has been well defined experimentally (Adams, et al. 1969; Baker, et al. 1988). It consists of two chains, A and B, linked by two interchain disulfide bonds (A7–B7 and A20–B19) and one intrachain disulfide bond within the A chain (A6–A11). This architecture forms a characteristic three-helix bundle comprising two α-helices from the A chain (yellow) and one α-helix from the B chain (blue), which together constitute the conserved structural core of insulin family members. In vertebrates, this disulfide scaffold is highly conserved and found in insulins, relaxins, and IGFs, although IGFs retain the C-peptide in their mature sequences. Sequences with the class I cysteine pattern have also been identified in several invertebrate phyla, including molluscs, annelids, and arthropods (Wu and Brown 2006; Veenstra 2010; Aslam, et al. 2011; Mizoguchi and Okamoto 2013; Cherif-Feildel, et al. 2019). Consistent with the high degree of conservation of this scaffold in vertebrates, all alternative cysteine-pattern classes identified in our dataset were found exclusively in invertebrate lineages, with the exception of Class II, which was also identified in amphibians.

Class II retains the cysteines corresponding to all three canonical disulfide bonds of Class I but contains an additional pair of cysteines positioned near the N-termini of the B- and A-chain regions. This sequence arrangement is consistent with the formation of an additional interchain disulfide bond, which, if formed as predicted, would increase cross-linking and constrain the relative orientation of the chain termini. Because only one cysteine pair remains unpaired beyond the canonical bonds, structure-prediction tools such as AlphaFold3 readily infer this additional connectivity. Class II represents a common cysteine framework in molluscs, annelids, and arthropods (Wu and Brown 2006; Veenstra 2010; Aslam, et al. 2011; Mizoguchi and Okamoto 2013; Cherif-Feildel, et al. 2019), and was also identified in the bryozoan *Watersipora subatra* and the vertebrate *Hyperolius riggenbachi*, a species of frog (Supporting File 5).

Together, Classes I and II comprise the vast majority of sequences in our dataset. Of the 6,604 sequences predicted by SignalP to contain a signal peptide (Teufel, et al. 2022), 5,618 (85%) were assigned to these two classes: 5,434 to Class I and 184 to Class II. By contrast, each of the other ten classes contained relatively few sequences and, with limited exceptions, remains largely undescribed.

Class III differs from the canonical Class I cysteine framework by lacking the pair of cysteines corresponding to the canonical intrachain disulfide bond of the A chain while containing an additional pair of cysteines positioned between the A- and B-chain regions. This arrangement is consistent with an alternative pattern of interchain connectivity. This architecture corresponds to the type α scaffold previously described in the nematode *C. elegans* (Pierce, et al. 2001). Class III sequences were recovered exclusively from other nematodes, including *Caenorhabditis brenneri*, *Caenorhabditis nigoni*, and *Caenorhabditis briggsae*.

Class IV shares features of the alternative cysteine framework observed in Class III but retains the cysteine pair corresponding to the canonical intrachain disulfide bond of the A chain. In addition, it contains an extra pair of cysteines near the N-terminal regions of the A- and B-chain regions, resulting in a framework containing four cysteine pairs in addition to the positional features associated with the canonical Class I insulin architecture. This pattern is consistent with the type β scaffold previously reported from *C. elegans* (Pierce, et al. 2001). Class IV sequences were recovered primarily from nematodes, including *Caenorhabditis brenneri*, *Caenorhabditis remanei*, *Cylicocyclus nassatus*, and *Ancylostoma ceylanicum*, with one additional sequence identified in the hemichordate *Saccoglossus kowalevskii*.

Class V retains the canonical Class I cysteine framework and contains an additional pair of cysteines positioned near the N-terminus of the B chain and the C-terminal region of the A chain. This arrangement introduces a potential disulfide bond between distant regions of the molecule that is not observed in other Classes. Relative to the canonical Class I framework, these peptides also contain additional residues within the intrachain disulfide loop of the A chain. The predicted structures are consistent with substantial modification of the canonical insulin architecture. Class V sequences were recovered from several molluscs and arthropods.

Class VI is defined by two sequences recovered from the ridgetail prawn, *Palaemon carinicauda*, which share high similarity with androgenic gland-specific insulin-like peptides (IAGs). IAGs are produced by the male-specific androgenic gland of crustaceans and function as key regulators of male sexual differentiation, reproductive development, and the maintenance of male phenotypes (Manor, et al. 2007; Ventura, et al. 2011). Whereas typical IAGs exhibit the canonical Class I insulin scaffold, the *P. carinicauda* sequences contain four additional cysteines, predicted to result in a rare five-disulfide architecture. One additional cysteine pair is positioned near the N-terminus of the B chain and the C-terminal region of the A chain, resembling the framework observed in Class V. The second additional pair is predicted to link the N-terminal region of the B chain to a region of the A chain immediately downstream of the cysteines corresponding to the canonical intrachain A-chain framework, resembling the pattern observed in Class IX. Thus, Class VI combines sequence-level features characteristic of Classes V and IX and represents a particularly elaborate cysteine framework within the insulin superfamily.

Class VII retains the canonical Class I cysteine framework and contains an additional pair of cysteines near the C-terminus of the B chain. In the predicted structures, this region adopts a bent or short helical conformation that brings the additional cysteines into proximity and is consistent with the formation of a local disulfide loop. Class VII sequences were recovered from two cnidarian species, the common jellyfish *Aurelia aurita* and the stony coral *Pocillopora damicornis*, suggesting that this unusual cysteine framework may represent a cnidarian-specific diversification of insulin-like peptides.

Class VIII sequences were recovered from three species of parasitic mites. These sequences retain the canonical Class I cysteine framework but contain an additional pair of cysteines near the N-terminus of the A chain. The predicted structures suggest that this additional cysteine pair may form an intrachain disulfide bond, resulting in an A chain with two intrachain disulfide bonds rather than the single intrachain bond characteristic of canonical Class I insulins.

Class IX retains the canonical Class I cysteine framework and contains an additional pair of cysteines positioned near the N-terminus of the B chain and in a region of the A chain immediately downstream of the cysteines corresponding to the canonical intrachain A-chain framework. This additional cysteine pair is predicted to form an additional cross-link not observed in any other insulins. Class IX sequences were recovered from Placozoa, one of the earliest-diverging non-bilaterian animal phyla.

Class X retains the canonical cysteine framework of Class I and contains an additional pair of cysteines near the N-terminus of the B chain, within an unusual N-terminal B-chain extension. The positioning of these additional cysteines is consistent with the potential formation of a small disulfide loop in the predicted structures. Whether this extended region containing the disulfide loop remains attached to the mature insulin-like peptide or is proteolytically cleaved to generate an independent bioactive peptide remains unresolved. Class X sequences were identified in the stony coral *Porites lutea*, the anemone *Nematostella vectensis*, and several other cnidarians.

Class XI sequences were recovered from Rotifera, a phylum of microscopic aquatic bilaterian animals. These sequences contain an insulin-like domain that retains the overall canonical Class I cysteine framework, together with an N-terminal extension of the B-chain region containing several additional cysteines. This extended region therefore represents a distinct and substantially more complex cysteine framework than those observed in the canonical insulin-like domain. Predicted structures suggest that the cysteine-rich extension may form a separate structured region containing several β-sheets and that it may be separated from the insulin-like domain by a predicted cleavage site. However, both the detailed three-dimensional organization of this region and its processing remain predictions. The sequence architecture nevertheless supports the presence of an additional cysteine-rich protein module of unknown function encoded within the same precursor.

Class XII retains the canonical Class I cysteine framework but contains one or two additional pairs of cysteines within the C-peptide region. The predicted structures suggest that these additional cysteine pairs may form intrachain disulfide bonds within the C peptide, potentially influencing precursor folding, processing, or give rise to a biologically active C-peptide-derived product. Unlike the representative structures for other classes, the pLDDT score for the Class XII representative shown in Figure 5 is comparatively low, as the C-peptide region, which is typically unstructured and excluded from confidence calculations in other classes, could not be excluded here without omitting the additional cysteines under investigation. Class XII sequences were identified in the insect *Drosophila virilis* and the tunicate *Styela clava*, with one and two predicted disulfide bonds within the C-peptide region, respectively.

### Coevolutionary analysis uncovers multiple structural segments in insulin-family

Coevolutionary analysis identifies pairs of amino-acid positions that tend to evolve in a coordinated manner, often because they participate in the same structural interaction or functional module. This approach can therefore reveal residues that are physically close in three-dimensional space or that share conserved functional roles. Because such inference requires large and diverse sequence alignments, the expanded dataset generated by our pipeline provided sufficient depth to perform a Direct Coupling Analysis (DCA) (Hopf, et al. 2019) of the insulin family. To quantify evolutionary constraints across the superfamily, we constructed a high-quality multiple-sequence alignment from the curated set of 7,020 sequences and applied DCA to infer residue–residue evolutionary couplings (Supporting File 6). We then compared these inferred couplings directly with the structural contact map of human insulin, allowing us to interpret coevolutionary signals within the context of known three-dimensional interactions.

The resulting coevolution matrix (**Figure 6A**) displays coupling strength as a color gradient, with darker cells indicating stronger evolutionary correlations. The upper triangle of the figure shows the computationally calculated contact map of human insulin in R conformation (PDB: 1AI0), a state in which the N-terminal region of the B chain adopts an extended α-helical structure, whereas the lower triangle shows inferred evolutionary couplings, enabling direct comparison between evolutionarily predicted residue–residue interactions and contacts observed in the known insulin structure.

**Figure 6.**
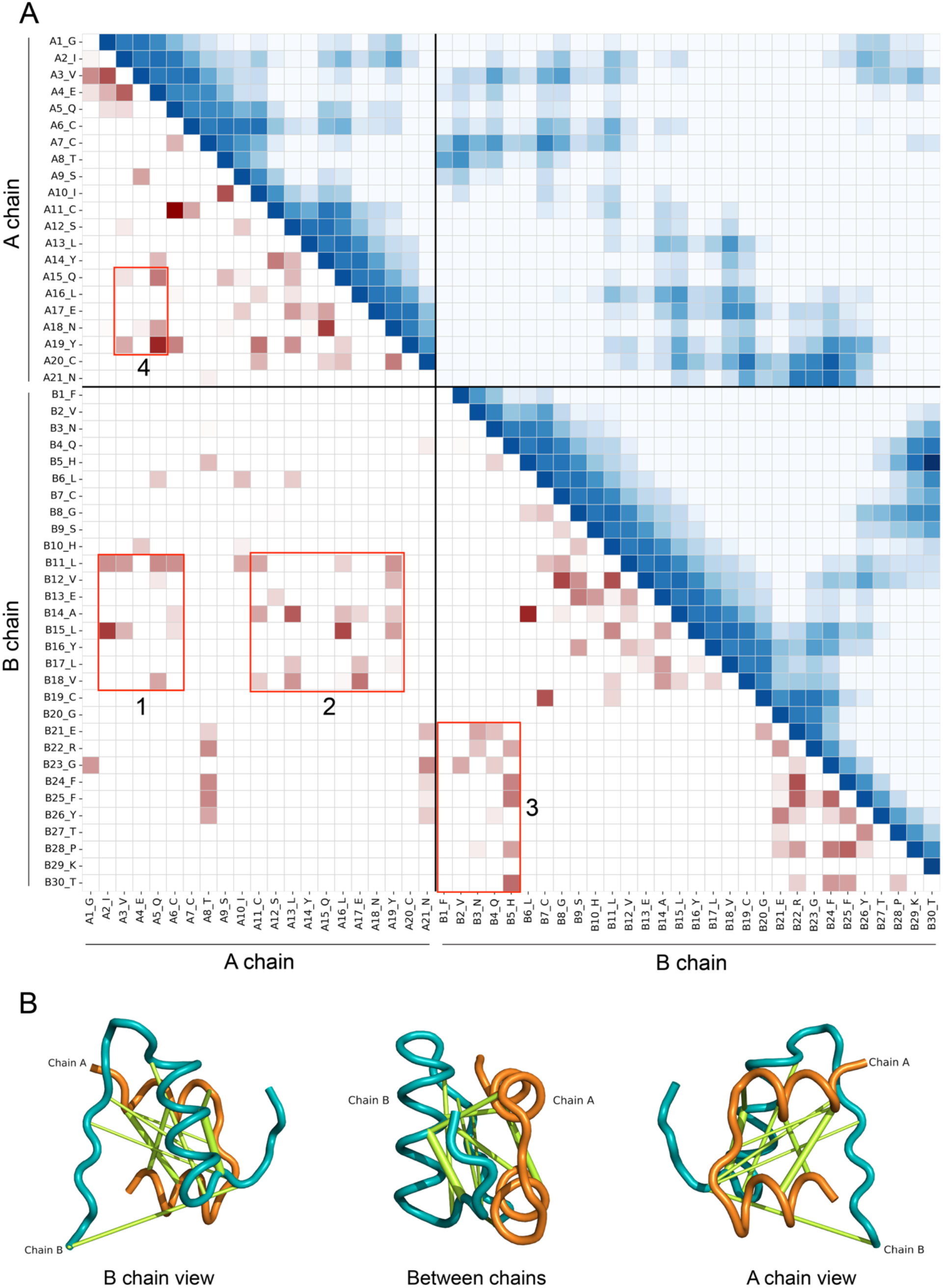
Comparison of structural contacts in human insulin with evolutionary couplings inferred from the expanded insulin-family dataset. (A) The upper triangle shows the Cα–Cα contact map of human insulin in the R-state conformation (PDB: 1AI0), with darker blue indicating shorter inter-residue distances. The lower triangle displays residue–residue evolutionary couplings inferred by Direct Coupling Analysis (DCA), ranging from weak (white) to strong (dark red). Mirrored signals near the diagonal identify three major coevolving structural segments corresponding to the A-chain helices, the central B-chain helix, and the B-chain C-terminal extension. Four prominent long-range coupling regions are highlighted. Regions 1 and 2 represent inter-chain couplings between the B-chain helical region and residues A1–A5 and A15–A20, respectively, indicating coordinated evolution of the B-chain helix with both A-chain helices. Region 3 represents intramolecular coupling between the N- and C-terminal regions of the B chain, consistent with their spatial proximity in the insulin structure. Region 4 highlights coupling between the two A-chain helices, reflecting structural constraints that maintain A-chain stability. Corresponding mirrored signals in the structural contact map demonstrate that these evolutionarily coupled residue pairs are also spatially proximal. (B) Three-dimensional representation of the identified coevolutionary interactions mapped onto the human insulin structure, with line thickness proportional to DCA coupling strength. Together, the correspondence between evolutionary couplings and structural contacts highlights conserved interactions that constrain the architecture of the insulin fold.

When strong signals appear mirrored across the diagonal of the matrix (**Figure 6A**), they indicate residue pairs that remain in close spatial proximity, such as those within the B-chain helix, and therefore coevolve to maintain structural integrity and function. Based on these mirrored patterns, the coevolution patterns revealed three major coevolving segments within the insulin molecule, the A-chain helices, the central helix of the B chain, and the C-terminal extension of the B chain. Beyond these short-range interactions, several long-range couplings were also evident. Strong inter-chain signals linked residues in the B-chain helical region with A1– A5 and A15–A20 (regions 1 and 2), indicating that both A-chain helices coevolve with the B-chain helix to preserve proper chain packing and receptor-relevant architecture. We further observed a pronounced intramolecular coupling between the N- and C-terminal regions of the B chain (region 3), consistent with the closed conformation of insulin. Finally, region 4 highlights coordinated evolution between the two A-chain helices themselves, reflecting structural constraints that maintain A-chain stability. A corresponding set of mirrored signals for regions 1–4 is also visible on the opposite side of the diagonal in the insulin contact map, further confirming that these residue pairs are spatially proximate and structurally constrained. Coevolutionary signals are visualized in **Figure 6B** (Supporting File 6; PSE file), which shows a 3D representation of these coevolutionary interactions in PyMOL, with line thickness proportional to coupling strength. Together, these coevolutionary signals underscore the conserved architecture of the insulin fold and reveal conformational determinants that have shaped its evolution.

## Discussion

Our survey provides a comprehensive picture of when and how the insulin signaling system arose in animals (**Figure 1**). The apparent absence of insulin-family ligands in Ctenophora, combined with our failure to detect insulin receptor (IR) homologs in this phylum, is consistent with prior reports (Sachkova, et al. 2021; Hayakawa, et al. 2022) and supports a model in which the insulin signaling system postdates the divergence of ctenophores from the rest of Metazoa. The situation in Porifera is more difficult to reconcile. Consistent with previous reports (Skorokhod, et al. 1999), we recovered IR-like sequences from multiple sponge species, supported both by phylogenetic placement within the IR clade (**Figure 2**) and by genomic analysis, arguing against a contamination explanation for the receptor. Yet despite an extensive search across 220 sponge species and nearly 2,000 transcriptomic libraries, with both sequence and structure-based search methods, we found no convincing evidence for an endogenous poriferan ligand. The one previously reported sponge insulin-like peptide (Robitzki, et al. 1989) appears, on re-examination, to be a rodent contaminant. This creates an apparent paradox: a receptor without a ligand. Several non-mutually exclusive explanations are possible. First, an ancestral poriferan ligand may exist but be too divergent in sequence to detect using homology-based approaches, particularly if sponge ILPs evolved under different selective constraints than the ligands used as queries. Second, the receptor may have originally interacted with a distinct, non-insulin-family ligand, with the canonical ILP-IR pairing established only later in the cnidarian/bilaterian lineage. Third, an ancestral ligand may once have existed in Porifera and subsequently been lost, while the receptor was retained and potentially repurposed for another function. Distinguishing between these scenarios will require functional characterization of the sponge IR-like receptor (e.g., ligand-binding assays or orphan receptor deorphanization approaches), more comprehensive genomic sampling across additional sponge lineages, and alternative ligand search strategies that do not rely on structure or sequence conservation alone.

While evidence for an insulin ligand–receptor pair in Porifera remains inconclusive, this signaling system is clearly present in Cnidaria (Steele, et al. 1996; Anctil 2009). Subsequently, insulin-family peptides underwent extensive diversification throughout animal evolution, particularly among invertebrates. Our phylogenetic analyses show that protostome and deuterostome sequences are generally well separated with non-bilaterian sequences grouping within protostome sequences (**Figure 3**). CLANS and PLM-based analyses independently show that deuterostome sequences form relatively well-defined clusters corresponding to insulin, IGF, and relaxin, whereas protostome and non-bilaterian sequences are more heterogeneous (**Figure 4**), suggesting that the greater sequence diversity of invertebrate peptides represents a feature of the insulin superfamily rather than an artifact of phylogenetic reconstruction.

Consistent with the sequence diversification observed in protostomes and non-bilaterians, nearly all structural innovation we detected (10 of 12 cysteine-connectivity classes) occurs in these lineages, despite vertebrates being disproportionately well-sampled in our dataset. Several of these classes were each identified in only one or a handful of species, and lineages such as Placozoa, Xenacoelomorpha, and Porifera remain represented by only a small number of sequences in current public databases. Given that novel scaffolds continued to emerge even within these sparsely sampled groups, including a previously undescribed cross-linked architecture recovered from just nine placozoan sequences (Class IX, described below), it is likely that many invertebrate lineages harbor additional, as-yet-undiscovered structural classes that will only become apparent as genomic and transcriptomic sampling of these underrepresented taxa expands.

These classes include those likely to form additional interchain and intrachain disulfide bonds, altered chain lengths and connectivity, and potentially additional protein domains (**Figure 5**). Several classes are particularly noteworthy. Class II represents a commonly observed scaffold in invertebrates that is characterized by two additional cysteines that likely form an additional interchain disulfide bond near the N-termini of the B- and A-chain regions, supplementing the three canonical disulfide bonds of the Class I architecture. While this framework is widespread among molluscs, annelids, and arthropods (Wu and Brown 2006; Veenstra 2010; Aslam, et al. 2011; Mizoguchi and Okamoto 2013; Cherif-Feildel, et al. 2019) and was also identified in the bryozoan *Watersipora subatra* here, we also identified a Class II sequence in the tree frog *Hyperolius riggenbachi*. A BLASTp search revealed similar sequences in several other anuran species (e.g., *Rana kukunoris, Rana temporaria, Lithobates pipiens*) suggesting this scaffold is present more broadly within this amphibian lineage. This represents a rare departure from the canonical three-disulfide-bond architecture of vertebrate insulins. Class IX, recovered from Placozoa, retains the canonical Class I cysteine framework but contains an additional pair of cysteines positioned near the N-terminus of the B chain and in a region of the A chain immediately downstream of the residues forming the canonical intrachain A-chain bond, predicted to form an additional cross-link not observed in any other insulin-like peptides in our dataset. This extends the known structural diversification of the insulin superfamily to an early-branching animal phylum. Similarly, Class X, identified from several cnidarian species, the earliest-diverging animals with an established insulin-IR signaling pair, also exhibits an unusual cysteine arrangement with an N-terminal B-chain extension containing two cysteines that likely form a disulfide bond. Whether this extension is retained in the mature peptide or proteolytically removed remains unknown, but the structural diversification observed in both Placozoa and Cnidaria suggests that structural diversification of the insulin molecule began very early in metazoan evolution. Class VI, represented by two sequences from the ridgetail prawn *Palaemon carinicauda*, illustrates the extent of structural elaboration possible within a single, functionally characterized lineage-specific context. These sequences share similarity with IAGs, which are established regulators of male sexual differentiation and reproductive development in crustaceans (Manor, et al. 2007; Ventura, et al. 2011). The predicted five-disulfide architecture, combining structural features of both Class V and Class IX, represents the most elaborate cysteine framework recovered in our dataset and suggests that functional specialization in this system can be accompanied by substantial structural innovation.

Beyond its evolutionary significance, this resource may have practical value for protein engineering. Several natural insulin variants have already inspired the design of insulin analogs: monomeric guinea pig and cone-snail insulins as templates for fast-acting insulins (Zimmerman and Yip 1974; Engholm, et al. 2015; Safavi-Hemami, et al. 2015; Xiong, et al. 2020; Xiong, et al. 2022), avian insulins for altered receptor potency (Simon, et al. 1974; Muggeo, et al. 1979), and viral insulins as tools for probing receptor agonism and antagonism (Altindis, et al. 2018; Zhang, et al. 2021). The scaffolds described here substantially expand this design space. For example, classes containing additional disulfide bonds (II, V, VI, VIII, XII) may offer templates for engineering enhanced conformational stability, a long-standing goal for improving insulin shelf life and manufacturability, while classes lacking the canonical A-chain intrachain bond (III) could inform efforts to understand the minimal disulfide requirements for a functional insulin fold. More broadly, the sheer diversity captured here suggests that natural sequence space may still contains largely untapped solutions to the specific biophysical trade-offs that continue to challenge therapeutic insulin analog design.

Coevolutionary analysis provides a complementary perspective on this structural diversity (**Figure 6**). Despite extensive variation in sequence and disulfide connectivity, strong evolutionary couplings were detected among the principal structural elements of the insulin fold. The A-chain helices, central B-chain helix, and B-chain C-terminal region displayed prominent coevolutionary signals, while additional long-range couplings connected the A- and B-chain regions that form the structural core. These patterns suggest that insulin-family evolution is characterized by a balance between structural conservation and peripheral innovation. The precise sequence and cysteine arrangement can vary considerably, yet residues involved in maintaining the overall fold remain subject to strong evolutionary constraints.

Beyond this, the dataset itself provides an additional resource for future research. By compiling a curated, phylogenetically broad seed set of 7,020 sequences we provide a valuable starting point for future homology searches as sequencing of underrepresented lineages continues. We anticipate that as new genomes and transcriptomes become available, particularly from early branching metazoans, this resource can be iteratively re-queried to test and refine the evolutionary scenarios proposed here and to identify new sequences and potentially new structural scaffolds.

Several limitations of this study should be considered when interpreting these results. First, not every sequence recovered by our pipeline is necessarily a bona fide insulin-family member; despite multiple filtering steps (length filtering, keyword exclusion, clustering, and manual structural inspection), some annotation errors or highly divergent false positives may remain in the dataset, particularly among the less well supported and taxonomically sparse groups. Second, our structural classification relies on predicted rather than experimentally determined structures. ESMFold and AlphaFold3 are generally reliable for well-conserved folds such as the insulin core, but predictions of disulfide connectivity become considerably less certain when multiple cysteine residues are positioned in close spatial proximity, as is the case for several of the more unusual classes described here (e.g., Classes VI, VIII, and XII). These predictions should therefore be treated as hypotheses that require experimental confirmation (e.g., mass spectrometry-based disulfide mapping, NMR, or crystallography) before being used to guide downstream functional or engineering work. Third, we deliberately did not attempt to predict proteolytic processing sites or mature peptide boundaries in this study, given the current lack of reliable, broadly validated tools for cleavage-site prediction outside of well-studied model systems. As a result, the chain boundaries depicted throughout this work should be considered provisional. This is a particularly important limitation for classes with additional N- or C-terminal extensions (e.g., Classes X, XI, and XII), where whether these extensions remain attached to the mature hormone or are cleaved to yield independent bioactive peptides remains entirely unresolved. Addressing this gap represents an important direction for future work, and improving processing- and modification-site prediction, especially for non-mammalian and invertebrate precursors, is a natural next step for this line of research. Finally, our dataset inherits the taxonomic biases of public sequence databases: well-studied model organisms and economically important species are overrepresented, while many invertebrate phyla remain sparsely sampled, meaning that some of the apparent lineage-restriction of individual structural classes may reflect sampling gaps rather than true evolutionary exclusivity.

## Conclusions

Together, our results reveal an insulin superfamily that is considerably more evolutionarily and structurally diverse than suggested by the canonical vertebrate insulin scaffold. The distribution of insulin-like sequences and insulin receptors supports a complex early evolutionary history in which the receptor component may predate the recognizable insulin ligand, while the canonical ligand–receptor signaling system is clearly established by the emergence of Cnidaria. Across later animal evolution, insulin-family peptides underwent extensive diversification, particularly in protostomes. Our identification of 12 distinct cysteine-connectivity classes demonstrates that the insulin fold represents an evolvable structural framework capable of accommodating additional interchain bonds, intrachain loops, altered chain connectivity, and potentially independent protein domains. The combination of this structural diversity with strong coevolutionary constraints highlights the balance between innovation and conservation that has allowed the insulin superfamily to persist across metazoan evolution. The resulting sequence and structural resource provide a foundation for future comparative and experimental studies of insulin-family evolution. As additional animal genomes and transcriptomes become available, this framework can be expanded to identify new sequence and structural classes. More broadly, the natural diversity captured in this dataset provides an extensive source of experimentally testable molecules for understanding insulin folding, receptor recognition, peptide processing, and potentially the design of next-generation insulin analogs.

## Materials and Methods

### Insulin Sequence Database Curation

We constructed a comprehensive insulin-family sequence database beginning with 243 seed sequences obtained from Feng et al. (Feng, et al. 2025), chosen for their validated coverage of all major insulin-family molecules across diverse metazoan lineages, including vertebrate relaxins, INSL peptides, and representatives of both deuterostome and protostome clades. This seed set was expanded using iterative PSI-BLAST searches against the NCBI non-redundant (NR) database (https://blast.ncbi.nlm.nih.gov). For each seed, we performed 3–5 PSI-BLAST(Altschul, et al. 1990) iterations using an E-value cutoff of 0.005 (Supporting Figure 1A). In each round, the updated set of hits was used to generate the PSSM for the subsequent iteration, and sequences were collected until reaching 5,000 total hits per seed or until the number of new sequences plateaued, indicating convergence. After pooling results from all seeds, the combined dataset contained ∼0.5 million sequences; removal of exact duplicates yielded 18,019 unique entries (Supporting Figure 1A-B; Supporting File 1).

To enrich for bona fide insulin-family precursors, we first applied a precursor-length filter, retaining only sequences 50–500 amino acids in length. This range corresponds to a pronounced peak of ∼15,000 sequences in our raw dataset (Supporting Figure 1B) and encompasses the typical 100–200 aa size of insulin-family precursors, while excluding sequences shorter than 50 aa that are overwhelmingly likely to be truncations given that signal peptides alone contribute 15–30 residues. At the upper end, the sharp decline in sequence counts beyond 500 aa further supports this cutoff: sequences longer than 500 aa are predominantly genome annotation-derived gene fusions or misannotated non-insulin proteins, rather than genuine insulin precursors. Next, to reduce redundancy while preserving true sequence diversity, we clustered the size-filtered dataset using CD-HIT (v4.8.1)(Huang, et al. 2010) at 95% identity, yielding ∼9,800 representative sequences (Supporting File 1). This threshold was selected after testing multiple identity cutoffs and identifying the point at which further reductions no longer meaningfully decreased redundancy while still preserving overall diversity.

To remove non-insulin contaminants, we implemented an iterative annotation-based filtering pipeline using both inclusion and exclusion keyword lists applied to sequence headers in the FASTA file. In each iteration, sequences whose annotations matched exclusion terms (e.g., “partial,” “receptor,” “zinc”) were removed, while those matching inclusion terms (e.g., insulin-family keywords) were retained. Remaining ambiguous entries were manually inspected, and any recurrent non-insulin keyword (for example, “Vitamin”) identified during this review was added to the exclusion list before rerunning the filter. This iterative refinement continued until no unrelated proteins remained. The final comprehensive inclusion and exclusion list used in this step is provided in Supporting File 7. After annotation-based filtering, approximately 7,200 sequences remained (Supporting File 1).

All remaining sequences underwent structure prediction using ESM-Fold (Lin, et al. 2023), a fast single-sequence structure-prediction model built on large protein language models (Supporting File 5). Unlike AlphaFold (Jumper and Hassabis 2022), which relies on multiple-sequence alignments and is computationally expensive, ESM-Fold generates structures directly from the input sequence and is substantially faster (∼1sec/seq), albeit with slightly lower accuracy; however, for single-chain proteins in the size range of insulin-family precursors, its accuracy is more than sufficient. Structure prediction was carried out with the pretrained esmfold_v1 model (esm Python package v1.0.3) on a single NVIDIA RTX 6000 GPU. Each sequence was processed individually using default parameters, and the resulting structure was written to a separate PDB file.

To further remove non-insulin-like entries, we sorted all predicted structures by file size and manually inspected those at both extremes: smaller files, which are most likely truncated proteins, and larger ones, which often correspond to misannotated non-insulin molecules. For each candidate, the predicted structure was aligned against human insulin scaffold and any sequence whose structure lacked an insulin-like backbone or did not contain an alignable insulin segment were removed. This process was continued until roughly 50 consecutive structures at each end of the distribution showed clear insulin-like folds, ensuring that no obvious non-insulin structures remained. Additionally, any sequences with anomalous predicted disulfide-bond counts, flagged by our disulfide-connectivity clustering pipeline (Methods; Structural framework classification), were individually examined by inspecting their predicted structures, aligning them against human insulin scaffold, and removing those that failed to show an insulin-like fold. The final curated dataset contained 6,969 high-confidence insulin-family sequences and served as the foundation for all downstream multiple-sequence alignment and evolutionary-coupling analyses.

We identified several phyla lacking insulin-family sequences, indicating that our method did not successfully recover insulin sequences from Placozoa, Phoronida, Platyhelminthes, Xenacoelomorpha, and Nemertea. We therefore searched for insulins in these phyla specifically using blastp against the nr database and tblastn against available transcriptome shotgun assemblies (TSAs) using the initial seed sequences as queries. Additionally, we searched available literature to identify any previous annotations (Nikitin 2015; Thiel, et al. 2021). We were able to retrieve insulin-family sequences from these phyla, which were added to the previously identified sequences, resulting in a final dataset of 7,020 sequences used for all subsequent analyses (Supporting File 1).

### Sequence Clustering

To visualize the global sequence similarity landscape of the insulin family, cluster analysis was performed using CLANS (Cluster Analysis of Sequences, version 29.05.2012). CLANS (Frickey and Lupas 2004) is a sequence-clustering tool that performs all-against-all BLASTp similarity searches using the BLOSUM62 matrix and uses a force-directed layout algorithm with iterative attraction–repulsion cycles to position sequences as points in two- or three-dimensional space based on their pairwise similarity scores. In this layout, BLASTp similarities create attractive forces between sequences shown as points, while unrelated sequences repel each other, causing clusters to form naturally. The complete curated dataset of 7,020 insulin sequences was used as input for all-against-all BLASTp comparisons with a stringent p-value threshold of 1×10⁻^19^ to identify high-confidence sequence relationships, and a p-value of 1×10⁻^7^ to show the connection lines between dots. (Supporting File 4). Using this p-value ensures that only highly similar sequences are placed within the same cluster. These thresholds were chosen empirically by testing multiple values and selecting those that provided optimal separation of known insulin-family groups (e.g., insulins, IGFs, relaxins) while maintaining meaningful connectivity patterns. The map can be loaded using the command and map files provided in Supporting File 4. The command to load the CLANS map is provided in Supporting File 7. Singleton sequences that did not cluster with any other sequences were hidden from the final visualization to emphasize major sequence groups and evolutionary subfamilies. Clusters were manually annotated using two complementary approaches. First, we assigned cluster identities based on known sequence annotations of representative members (e.g., human insulins, IGFs, and relaxins). Because many entries in our dataset carry automatically generated NCBI annotations, only densely populated regions corresponding to well-defined families, such as the relaxin family, could be reliably identified using this approach. Second, we annotated clusters using phylogenetic information. To do this, we downloaded the NCBI taxonomy names and nodes files (https://ftp.ncbi.nih.gov/pub/taxonomy/taxdump.tar.gz; Supporting File 4), loaded them into the CLANS interface, searched for the Metazoa NCBI TaxID, and assigned the resulting groups. FASTA files for all phylum-specific sequence sets used in this process are provided in Supporting File 4. To quantify the number of sequences and species represented in these FASTA files, we counted the total number of sequences and the number of unique species names (enclosed in square brackets) for each phylum and summarized these values in Supporting File 4. To generate Figure 4A, we colored higher-level taxonomic categories such as Protostomia and Deuterostomia, along with other minor groups shown in Figure 1.

### Cladogram creation

To characterize the taxonomic distribution of sequences in the curated insulin family dataset, we used the summary tables in the Excel file included in Supporting File 4 (described in the Sequence Clustering section above) as the basis for generating a phylum level cladogram. The Excel file reports the total number of sequences, the number of species represented for each phylum. The Newick tree was manually constructed based on the NCBI taxonomy hierarchy, with all outputs manually verified by the authors. This annotated NWK file was uploaded into iTOL (https://itol.embl.de), where we adjusted tree dimensions and added the sequence count, color, and label datasets.

### Porifera sequence data collection and processing

Publicly available sponge RNA-sequencing datasets were retrieved from the NCBI Sequence Read Archive (SRA) by restricting the taxonomic search to Porifera. The search was conducted in April 2026. Raw reads for each SRA were downloaded using sratoolkit.3.3.0(Information) 2026) with the command “fastq-dump --split-files”. Adapter and low-quality sequences were removed using Trimmomatic-0.39(Hyatt, et al. 2010) with the command “trimmomatic-0.39.jar PE -phred33 LEADING:3 TRAILING:3 SLIDINGWINDOW:4:15 MINLEN:50”. These settings removed bases with quality scores below 3 from the beginning and end of each read, trimmed reads when the mean quality within a four-base sliding window fell below 15 and discarded reads shorter than 50 nucleotides after trimming. Open reading frames were predicted and translated from each transcriptome assembly using Prodigal v2.6.3(Hyatt, et al. 2010). The resulting translated protein sequences were used as the transcriptome-derived proteome for downstream insulin-family searches.

Publicly available sponge genome assemblies were retrieved from the NCBI Assembly database using Porifera as the taxonomic filter in April 2026. The assembly for each genome was annotated using online AUGUSTUS (https://bioinf.uni-greifswald.de/webaugustus/)(Hoff and Stanke 2013). The predicted protein sequences were exported and used as the genome-derived proteomes in subsequent insulin-family and insulin-receptor searches.

### Searches for insulin-like proteins in Ctenophora and Porifera

Notably, no insulin-family sequences were detected in Ctenophora or Porifera using this pipeline. To ensure that this apparent absence was not an artifact of our search strategy, we performed an extensive series of additional analyses. Using human and Cnidarian insulin and insulin receptor sequences as queries, we searched Ctenophora and Porifera in both the NR and TSA databases on NCBI website (https://blast.ncbi.nlm.nih.gov). No insulin or insulin-receptor hits were identified in Ctenophora across either database. In contrast, although no insulin-like sequences were detected in Porifera, multiple insulin-receptor homologs were found (Robitzki, et al. 1989) (Supporting File 2). This unexpected discrepancy prompted a deeper investigation into the possible presence of Poriferan insulin-like molecules.

An in-house Porifera dataset comprising more than 200 species and 1,945 samples and approximately 160 million predicted protein sequences with 50–500 aa in length was mined. We performed BLAST (version 2.16.0) against the Porifera dataset using different insulin sequences as query existing in our insulin dataset, covering all classes of Figure 5. The top-scoring candidate hits were manually examined and retained only if they contained the characteristic cysteine-rich motif typical of insulin-family peptides. These verified hits were then aligned, and the conserved cysteine-rich region was extracted to build a hidden Markov model (HMM) (http://pfam.xfam.org). The resulting HMM profile was used to search the sponge transcriptome database for additional insulin-like sequences. Newly recovered hits were re-evaluated, aligned, and incorporated into an updated HMM, and this iterative search procedure was repeated four times to obtain the final set of insulin-like hits. As a result, only 129 insulin-like sequences were detected from 1,945 SRA samples. The sequences are available in Supporting File 1.

To evaluate whether putative insulin-like sequences represented conserved sponge genes, we examined their distribution across the in-house Porifera sponge transcriptomes dataset. Each SRA accession was classified as positive when it contained at least one insulin-like hit. SRA samples were grouped by species, and the total number of sampled SRA libraries, number of positive libraries, and proportion of positive libraries were calculated for each species.

Genomic support for the transcriptome-derived insulin-like sequences was evaluated using a panel of 56 sponge genome assemblies and their corresponding predicted proteomes. All 56 species were represented in the transcriptomic dataset by at least one SRA sample. Each transcriptome-derived insulin-like sequence was searched against the complete proteome dataset generated from these 56 genome assemblies. Using BLASTP v2.16.0 with an E-value cutoff of 1 × 10⁻¹⁰, no significant hits were detected. To account for the possibility that short genes may have been missed during genome annotation and gene prediction, we additionally performed direct tblastn searches against the genomic DNA sequences of all sponge genome assemblies. Consistent with the BLASTP results, no significant matches were identified in any genome assembly.

To test whether insulin-like sequences could be detected directly from genomic resources, the same insulin-mining pipeline used for the transcriptome collection was applied independently to each sponge genome assembly derived proteome and no hit was obtained.

As an alternative to sequence-based homology searches, we also used Foldseek v10 (van Kempen, et al. 2022) to search for proteins with insulin-like folds. Foldseek can search a sequence-based database using a structural query by inferring the structural characteristics of the target sequences and comparing them to the query, completely bypassing the need for explicit 3D structure prediction. Therefore, we extracted proteins of length 80-200 amino acids from the poriferan database and clustered them to 80% identity using easy-clust of MMseqs2 v18.8cc5c(Steinegger and Söding 2017) with (min-seq-id 0.80 -c 0.80 --cov-mode 0 --cluster-mode 0 -s 7.5) and used these to construct a custom database with foldseek createdb. We used foldseek easy-search using human insulin full length predicted structure with AlphaFold3 (Abramson, et al. 2024) as a query against this database with an e-value threshold of 1e-1. Again, no genuine insulin-like fold candidates were identified; all apparent hits were traceable to contaminating sequences from non-Poriferan species (Supporting file 1).

### Porifera Insulin Receptor exploration

Having found no evidence for insulin-like ligands by either sequence or structure, we asked whether the downstream signaling machinery might nevertheless be present in Porifera. Following the same pipeline used in the insulin-like sequence mining, BlastP searches for insulin receptor homologs was performed using our manually identified sponge insulin receptor sequences as query and against our in-house dataset revealed many hits across diverse Poriferan lineages, with 187 sequences having confirmed genomic location (Supporting File 1), consistent with previous reports (Robitzki, et al. 1989).

### Receptor tyrosine kinase tree construction

We constructed a reference dataset comprising all human RTKs as defined by Table 1 in Filis et al. paper (Filis, et al. 2023), together with 187 putative insulin receptor sequences identified from our in-house poriferan database. Phylogenetic analysis was performed as described in Methods (Insulin Tree Construction). Briefly, sequences were aligned using MAFFT, and the resulting multiple sequence alignment was trimmed with trimAl using a gap threshold of 0.5. ModelFinder identified Q.mammal as the best-fitting substitution model, which was subsequently used for phylogenetic tree inference. The resulting tree was visualized using iTOL and is shown in Figure 2A.

To further validate this classification, we curated a broader dataset of insulin receptors spanning the major animal phyla represented in Figure 1. We first used the human insulin receptor as a BLASTp query against the NCBI nonredundant (NR) database to identify insulin receptor sequences from each phylum. Three representative sequences from each of the ALK, LTK, and ROS1 families were included as the closest identified outgroup. The alignment was filtered with trimAl to retain positions present in at least 50% of sequences, then used to infer a maximum-likelihood phylogenetic tree in IQ-TREE2(Nguyen, et al. 2015), which selected Q.mammal as the best-fit evolutionary model and calculated robust branch support. (Methods: Insulin tree construction). (Figure 2B; Supporting File 2).

### Group 1 and Group 2 phylogenetic analysis

To test whether the two divergent protostome clades identified in the full-dataset phylogeny (Groups 1 and 2; Figure 3A) represent relaxin-like peptides, we constructed a focused maximum-likelihood tree containing these sequences and representative members of the major insulin-superfamily subgroups. Reference sequences were obtained through BLASTp searches against the NCBI nonredundant (NR) database. We retrieved the top ten hits for *Drosophila melanogaster* Dilp8, a previously characterized protostome relaxin (Vallejo, et al. 2015), and the top ten hits for each of human insulin, IGF, and relaxin, five from Deuterostomes, and five from Protostomes. These sequences were combined with all Group 1 and Group 2 sequences from the curated dataset (Supporting File 3). Sequences were aligned using MAFFT v7.407, After removing poorly represented alignment positions with trimAl, we reconstructed a maximum-likelihood phylogeny in IQ-TREE v2.4.0 (Nguyen, et al. 2015), ModelFinder selected Q.mammal as the best-fit amino acid substitution model under the Bayesian information criterion. The resulting tree was midpoint-rooted and visualized using iTOL (Interactive Tree Of Life, https://itol.embl.de) (Methods: Insulin tree construction). All input sequences, alignments, and tree files are provided in Supporting File 3.

### Insulin tree construction

A maximum-likelihood phylogenetic tree was constructed from the curated 7,020 insulin-family sequence dataset. The protein sequences were aligned with mafft v7.407 (Katoh, et al. 2005), and a maximum likelihood tree was reconstructed using IQ-TREE v2.4.0 (Nguyen, et al. 2015). The model of evolution was calculated to be Q.plant by the Bayesian information criterion with ModelFinder (Kalyaanamoorthy, et al. 2017), we restricted ModelFinder to the general matrices (LG, WAG, JTT) plus lineage-specific Q matrices to keep run time manageable. Bootstrap values were calculated with 1000 replicates with UFBoot (Hoang, et al. 2018). Branch support was assessed using 1000 replicates of the SH-like approximate likelihood ratio test (SH-aLRT) alongside 1000 replicates of the Ultrafast Bootstrap approximation (UFBoot2) optimized with a regularized stubborn tree search (-bnni) to mitigate severe model violations. Tree search heuristics were constrained to a minimum of 10 initial candidate trees (-ninit 10) with the stopping criterion set to 200 unsuccessful iterations (-nstop 200). The IQtree2 command is available in Supporting file 7.

The final tree was uploaded to the iTOL (https://itol.embl.de) web server and visualized as an unrooted tree. To distinguish the different sequence groups, annotation datasets were generated using the custom scripts provided in Supporting File 7, with the FASTA files for animal groups (Supporting File 1) and molecular families (Supporting File 4) as inputs. The resulting annotation datasets were then uploaded to iTOL to color-code the corresponding clades.

To further examine the internal diversity of the three deuterostome insulin superfamily groups, we constructed a phylogenetic tree containing only deuterostome insulin, IGF, and relaxin sequences (Figure 3C). This tree was generated using the same pipeline described above for the full dataset. ModelFinder Plus selected Q.mammal as the best-fit amino acid substitution model for this subset.

### Protein language model-based clustering

To complement traditional phylogenetic approaches and visualize sequence relationships through an independent method, we employed the Protein Language Model ESM-2 v2.0.0 for embedding-based clustering (Lin, et al. 2023). For each protein, ESM-2 produces a per-residue embedding matrix, where one axis represents sequence positions and the other represents learned features as a list of numbers. To obtain a single fixed-length representation per protein regardless of sequence length, embeddings were averaged across the residue axis, yielding a mean-pooled vector summarizing the biochemical character of the entire sequence.

The mean-pooled embedding vectors were standardized feature-wise by z-score normalization using the StandardScaler implementation in scikit-learn(Hoff and Stanke 2013). The standardized vectors were then projected into two dimensions using the Python UMAP-learn implementation of UMAP (Uniform Manifold Approximation and Projection), with n_neighbors=10, min_dist=0.1, n_components=2, and random_state=42. These parameters were selected to generate a reproducible low-dimensional representation emphasizing local relationships among sequences. Density-based clustering was subsequently applied to the two-dimensional UMAP coordinates using the DBSCAN implementation in scikit-learn, with eps=0.6 and min_samples=10’. DBSCAN assigned sequences in sufficiently dense regions to clusters and classified points outside these regions as noise; cluster identifiers were shifted by one so that noise points were labeled 0 and clusters were numbered beginning with 1.

To interpret the resulting embedding space from two complementary perspectives, points were colored in two ways: first, according to broad phylogenetic affiliation of Protostomes and Deuterostomes (As in Figure 3A) based on the taxonomic group files defined during CLANS-based sequence clustering (see Methods: Sequence Clustering above); and second, according to molecular family (Insulin, IGF, and Relaxin, Supporting Figure 4) as defined by the deuterostome subfamily groupings established in the same CLANS analysis (Figure 4A), to assess whether Protein Language Model (PLM)-derived clusters reflect functional and structural subfamily identity independently of taxonomic origin. Both coloring schemes are shown in an overlaid representation in Figure 4B. The clustering files are available in Supporting File 4, and codes are available in Supporting File 7.

### Structural framework classification

We predicted signal peptides for all 7,020 sequences using SignalP v6.0 (Teufel, et al. 2022) in slow-sequential mode (command in Supporting File 7). Signal peptides were identified in 6,007 sequences; the remaining 1,013 lacked a detectable signal sequence (Supporting File 5).

Many sequences without predicted signal peptides appeared N-terminally extended or truncated, likely due to genome-based prediction errors, sequencing artifacts, or incomplete transcript assemblies. Because protein-coding mRNAs typically begin with methionine and alternative translational start sites are common, we reasoned that N-terminally truncated variants might still yield valid signal-peptide predictions. To address this, we wrote a custom Python script (Supporting File 7) that generates all subsequences beginning at each internal methionine (Supporting File 5).

Re-running SignalP 6.0 on the sequence subsets and replacing the longest variant containing a predicted signal peptide with the original sequence recovered 597 additional signal-peptide-containing sequences. This increased the total to 6,604 sequences with predicted signal peptides; 416 sequences were excluded for lacking any identifiable signal-peptide form (Supporting File 5).

Signal peptides were trimmed and mature sequences were aligned with MAFFT v7.407 (Supporting File 5). Because raw sequence coordinates are not directly comparable across divergent sequences, we mapped each cysteine to its corresponding column in the multiple-sequence alignment (MSA) using a custom Python script (Supporting File 7). This identified homologous cysteines. An overview of this procedure is illustrated in Supporting Figure 5. In this workflow, each sequence is first scanned to identify the positions of all cysteine residues in its native, unaligned form (Supporting Figure 5A–B). After generating a global MSA, the script then remaps each cysteine to its aligned position in MSA and converts sequence cysteine positions to cysteine positions in MSA (Supporting Figure 5C). Once all sequences were converted to MSA-indexed cysteine patterns, we collapsed sequences sharing identical cysteine-position sets (Supporting Figure 5D). This step removes redundant connectivity patterns and reduces the dataset to a non-redundant set of unique cysteine-organization types. These unique patterns represent distinct potential disulfide-bonding architectures. The CSV files of sequences before and after disulfide connectivity deduplication are available in Supporting File 5. The resulting reduced set was then manually inspected to identify unusual, rare, or non-canonical cysteine arrangements that might represent novel structural classes. In total, we identified 12 structural groups. Detailed information for each group, including species composition, sequence counts, and class assignments, is provided in Supporting File 5, along with a PSE file visualizing each group’s representative structure with disulfide bonds rendered explicitly.

### Coevolutionary analysis

To infer evolutionary couplings, we used the EVcouplings v0.2.1 (Hopf, et al. 2019) framework monomer protocol Supporting File 7), which implements DCA through the plmc pseudo-likelihood maximization model and includes optional scoring modules that help distinguish statistically meaningful couplings from background noise. Because our study already assembled a comprehensive insulin-family sequence dataset, we constructed the MSA directly from these sequences using MAFFT v7.407. This curated alignment was then provided as the input for the EVcouplings monomer pipeline for downstream coupling inference.

In the first stage of the EVcouplings monomer pipeline, EVcouplings applies sequence reweighting to correct for uneven sampling across taxa. Reweighting is controlled by θ (theta), the sequence-identity threshold used to group closely related sequences into a single representative. Values between 0.8 and 0.9 are commonly used; we tested both and observed no substantial differences in downstream coupling patterns, and therefore selected θ = 0.85. Under this setting, sequences sharing >85% identity are down-weighted, preventing phylogenetically clustered sequences from dominating the evolutionary signal.

To ensure alignment quality, the module enforces both sequence-level and column-level completeness thresholds. Sequences with <30% alignment coverage (minimum_sequence_coverage = 30) are removed to eliminate partial or poorly aligned entries, while alignment columns present in fewer than 20% of sequences (minimum_column_coverage = 20) are excluded to reduce noise from sparsely populated sites. These thresholds were selected empirically: starting with stricter cutoffs, we gradually lowered them and confirmed that including additional sequences did not degrade the evolutionary signal or introduce noise, consistent with our dataset already being highly filtered.

After filtering and reweighting, EVcouplings reports the effective sequence number (N_eff), which quantifies the amount of usable evolutionary information in the alignment. For our dataset, the final alignment contained 7,020 sequences, of which 96 of 110 positions (87.3%) passed column-coverage filtering, and reweighting at θ = 0.85 yielded an N_eff of 2,348.588, providing sufficient statistical depth for reliable estimation of residue– residue evolutionary couplings. This quality-controlled MSA was then passed to the couplings stage for DCA inference.

In the second stage of the pipeline, the couplings module performs DCA to infer pairwise evolutionary couplings between alignment positions. We used the standard protocol, which calls the plmc pseudo-likelihood maximization framework to estimate the parameters of an undirected graphical model from the filtered MSA. Model optimization was run for 500 iterations, a commonly used setting that ensures convergence while minimizing overfitting. To prevent overfitting, we applied L2 regularization using standard values of λ_J = 0.01 (scaled by sequence length) and λ_h = 0.01. A minimum sequence separation of six residues (min_sequence_distance = 6) was enforced to distinguish trivial short-range contacts from meaningful long-range interactions. Final coupling strengths were converted into predicted contact scores using the logistic-regression scoring model implemented in the EVcouplings framework (Supporting File 6/couplings/INS85_CouplingScores).

To interpret the biological meaning of the inferred evolutionary couplings, it is essential to compare them directly with a structural contact map. DCA identifies residue pairs that co-evolve because they are structurally or functionally related, while the contact map provides the three-dimensional framework needed to determine which couplings reflect true spatial proximity or functional relationships. To enable this comparison, we generated a unified two-dimensional heatmap that contrasts structural contacts and evolutionary couplings in a single matrix (Figure 6A). The upper triangle contains the structural contact map derived from the insulin R-state structure (PDB 1AI0). Residue identities for chains A and B were extracted and labeled by chain, residue number, and one-letter amino-acid code (e.g., B25_F). Pairwise Cα distances were calculated, and a 13 Å cutoff was applied to retain biologically meaningful contacts associated with helix–helix packing and inter-chain interactions. Distances were then linearly normalized so that shorter distances map to higher contact intensities. The lower triangle displays evolutionary couplings derived from DCA. EVcouplings scores (Supporting File 6/couplings/INS85_CouplingScores), which quantify the likelihood that two residues co-evolve, were independently log-normalized and scaled to the 0–1 range, ensuring that their dynamic range is comparable to the normalized structural distances in the upper triangle. To visualize co-evolutionary constraints on the human insulin structure, EVcouplings automatically produced a pml script for mapping pairwise evolutionary couplings onto a 3D model. We first predicted the structure of human insulin using AlphaFold3 (Abramson, et al. 2024) and loaded the resulting model into PyMOL (LLC. 2010). The EVcouplings-generated pml file was then applied to the model, after which we removed the signal-peptide and C-peptide regions to focus exclusively on the mature insulin fold. The final annotated structure was saved as a .pse file (Supporting File 6), and three representative orientations of this model are shown in Figure 6B.

## Data Availability

The dataset is available on figshare at https://doi.org/10.6084/m9.figshare.34105398.

## Acknowledgements

The support and resources from the Center for High-Performance Computing (CHPC) at the University of Utah are gratefully acknowledged. Structures were predicted under the University of Utah AlphaFold3 license. Claude AI was used to assist with code debugging. The authors reviewed all code and take full responsibility for the analyses.

