## Supporting Figures for "The origin and diversity of insulin sequences and structures in nature"

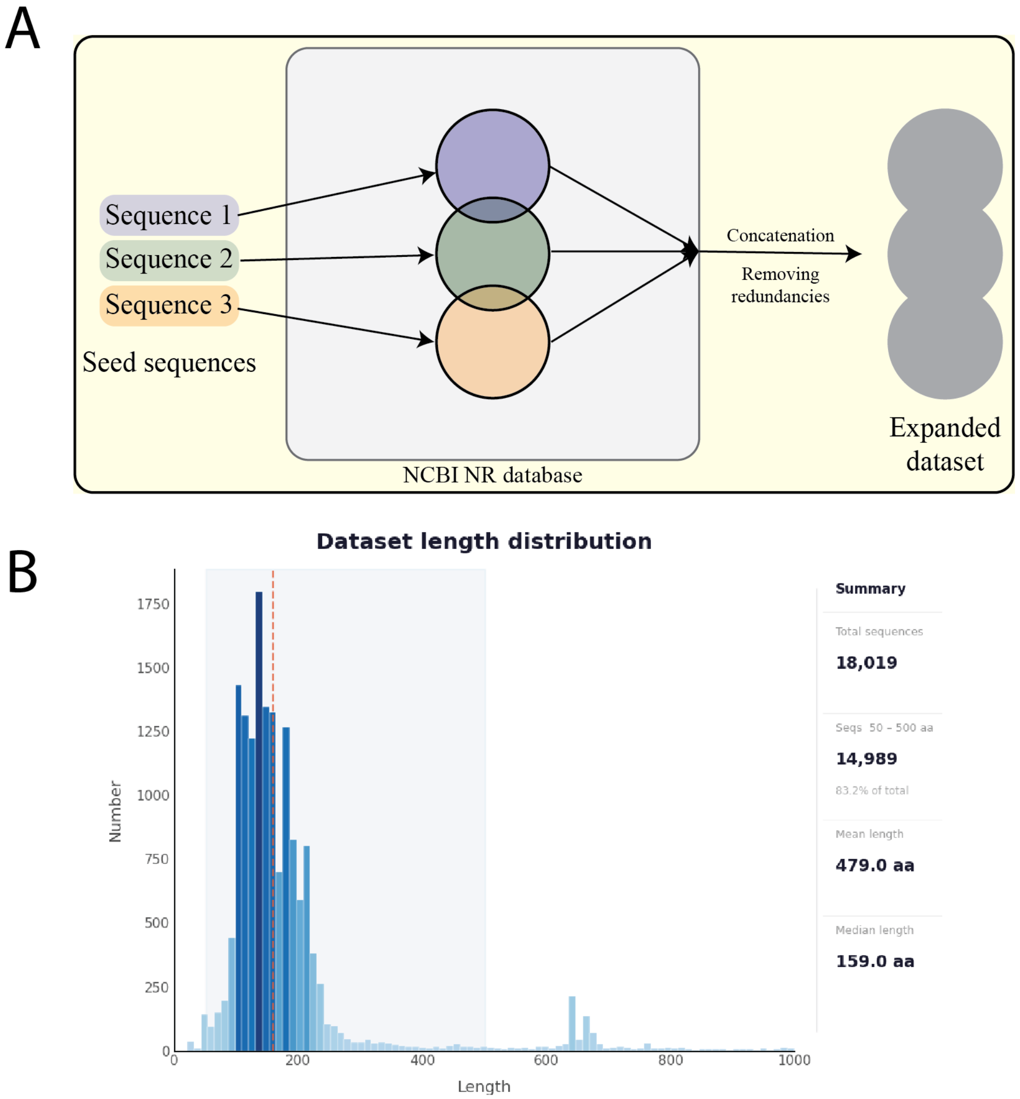


**Supporting Figure 1. Dataset curation workflow. (A)** The initial seed set of 243 validated insulin-family sequences was expanded through iterative PSI-BLAST searches against the NCBI non-redundant (NR) database, with each seed sequence queried for up to five iterations. In each iteration, the updated set of sequences was used to build the position-specific scoring matrix (PSSM) for the subsequent round. Although substantial overlap between seed hits was anticipated given their shared insulin-family origin, the subtle sequence differences between seeds ensured that each seed uniquely contributed new hits in the final dataset, progressively expanding the recovered sequence space without manual intervention. After pooling all hits across seeds, the combined dataset reached approximately 500,000 sequences. Removing exact duplicates reduced this to 18,019 unique insulin-like sequences. **(B)** Length distribution of retrieved sequences prior to curation. Approximately 15,000 sequences fall within the characteristic 50–500 amino acid range of insulin family precursors, forming the dominant peak used to guide downstream filtering.

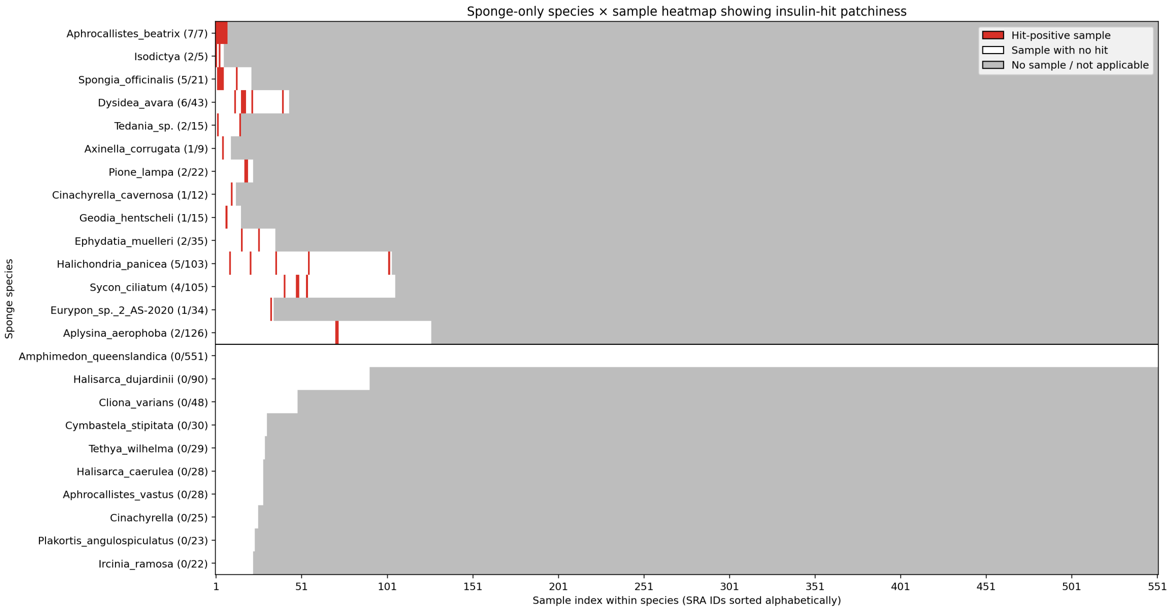


**Supporting Figure 2.** Species-by-sample heatmap of Porifera transcriptome libraries revealing a patchy within-species distribution, with hit-positive libraries scattered among predominantly negative libraries rather than being consistently recovered across conspecific transcriptomes. Species name is followed by the number of insulin hits and the number of interrogated transcriptomes in parenthesis.

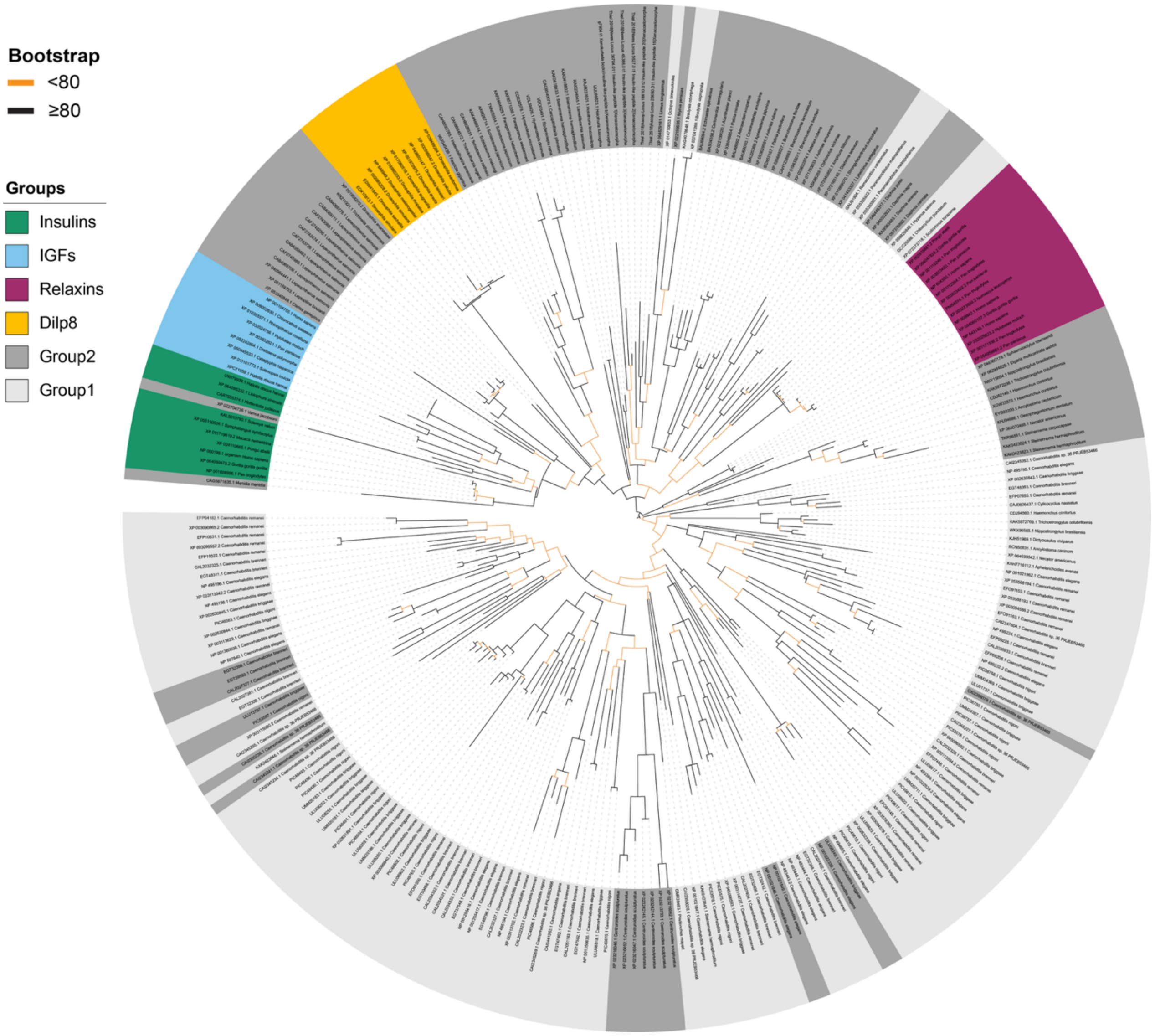


**Supporting Figure 3. Identity of Group 1 and Group 2 sequences.** We hypothesized that Group 1 and Group 2, the protostome sequences that cluster with deuterostome sequences in Figure 3A, represent protostome relaxins. To test this, we assembled a reference dataset of 10 insulins, 10 IGFs, 10 relaxins, and 10 Dilp8-like sequences (established protostome relaxins) and added the Group 1 and Group 2 sequences. The outer ring marks Insulins (green), IGFs (light blue), Relaxins (magenta), Dilp8-like sequences (yellow), Group 2 (dark gray), and Group 1 (light gray). Branch color indicates bootstrap support (black, ≥80; orange, <80). Insulins and IGFs form adjacent, compact clades, and relaxins form a distinct clade. Dilp8-like sequences are separated from the insulin/IGF cluster, and most Group 2 sequences are located near them. Most Group 1 sequences are placed close to the relaxin clade. These results are consistent with Group 1 and Group 2 being protostome relaxins, although the placement of Group 2 is closer to Dilp8-like sequences.

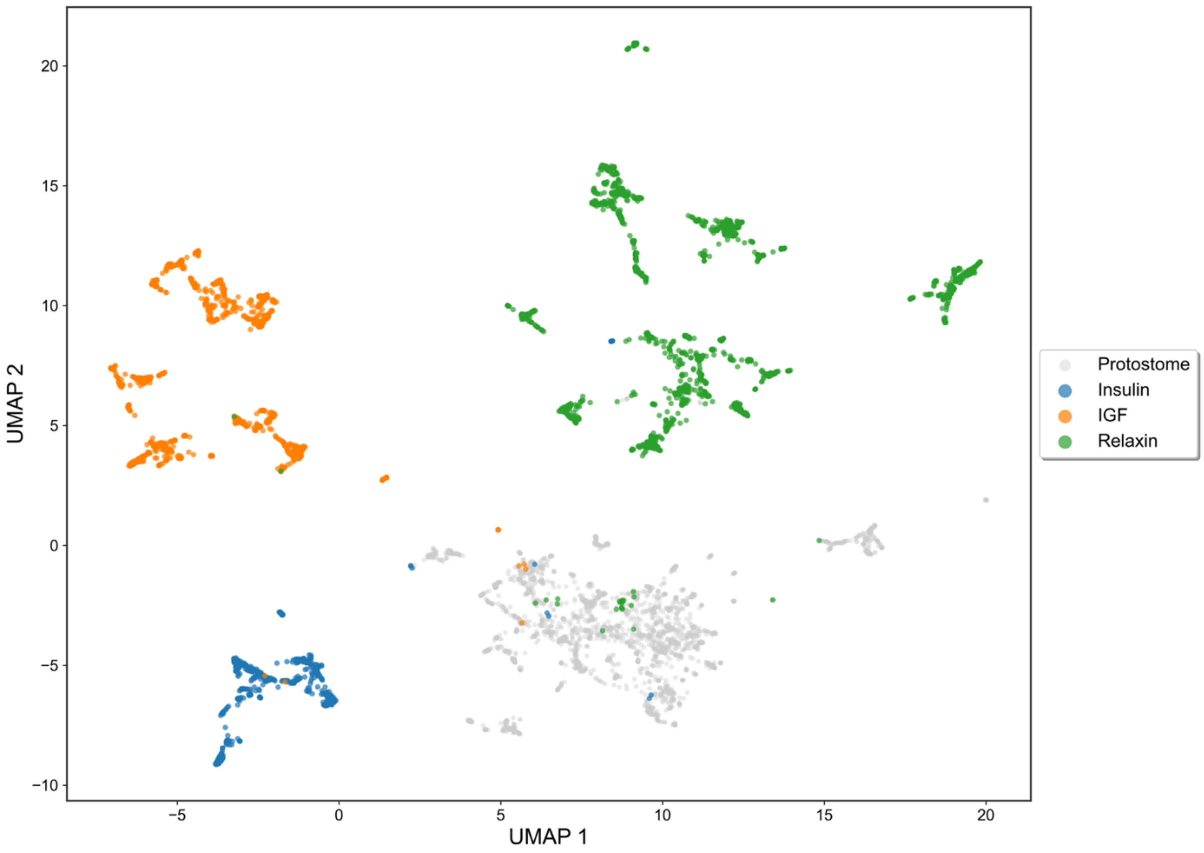


**Supporting Figure 4. UMAP embedding of ESM-2 protein representations reveals subfamily-level clustering within deuterostomes.** To assess whether the subfamily-level clustering observed in Figure 4A is also reflected in Figure 4B, we colored the ESM-2-based UMAP using the same insulin, relaxin, and IGF groupings identified in Figure 4A. Sequences are colored by group: protostomes (gray) and three deuterostome subfamilies, insulin (blue), IGF (orange), and relaxin (green). The three subfamilies form distinct clusters in the UMAP embedding, mirroring their separation in Figure 4A. Because this approach relies on learned sequence representations rather than explicit multiple sequence alignments or phylogenetic inference, it provides an independent line of evidence that subfamily-level divergence within the insulin superfamily is recoverable from sequence information alone.

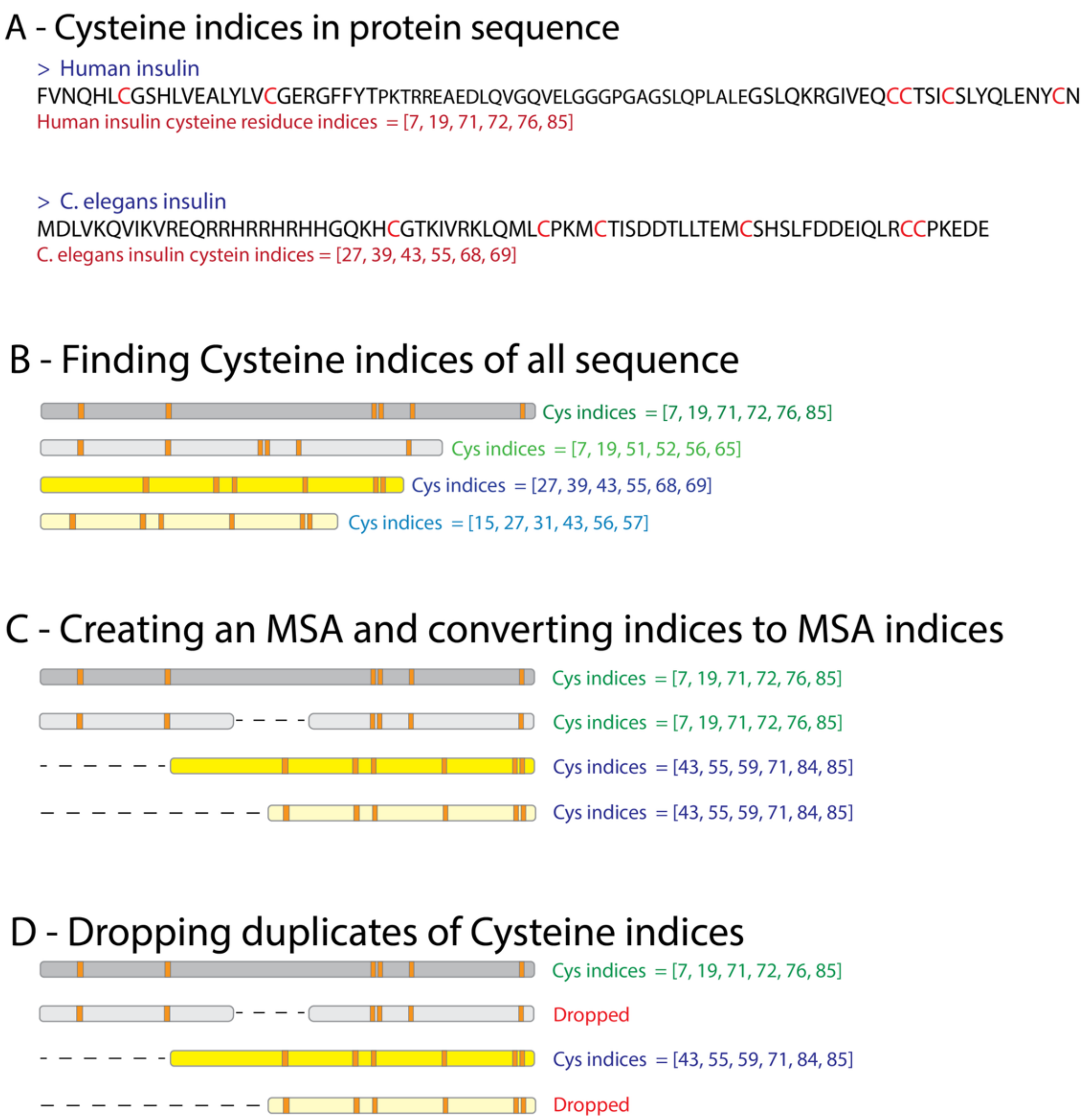


**Supporting Figure 5. Workflow for identifying cysteine-based structural outliers within the insulin-family dataset. (A)** Cysteine positions are listed for representative sequences, illustrated using human insulin and C. elegans insulin as examples. **(B)** Cysteine indices are computed across all sequences. Four schematic sequences are shown: the first represents human insulin; the second is a human insulin-like sequence with a shorter C-peptide; the third represents C. elegans insulin; and the fourth is a C. elegans-like sequence with a shorter N-terminal extension. Notably, even closely related sequences, shown as yellow and grey pairs, yield different cysteine position lists, making direct comparison based on raw indices unreliable. **(C)** Sequences are aligned using MAFFT, and raw cysteine indices are converted to MSA-aligned positions to ensure homologous residues are directly comparable across divergent sequences. After alignment, the cysteine position lists become consistent and identical for homologous proteins, resolving the ambiguity seen in panel B. **(D)** Redundant cysteine-position patterns are collapsed into a non-redundant set of connectivity classes, which are then used to identify rare or non-canonical disulfide-bond topologies in downstream structural analyses.
